# FGF and BMP pathway genes are expressed in segmentally iterated domains during segmentation of germ layers in a tardigrade

**DOI:** 10.1101/2024.01.29.577774

**Authors:** Kira L. Heikes, Lillian D. Papell, Noemi Gavino-Lopez, Bob Goldstein

**Affiliations:** Biology Department, University of North Carolina at Chapel Hill, Chapel Hill, NC, USA; Curriculum in Genetics and Molecular Biology, University of North Carolina at Chapel Hill, Chapel Hill, NC, USA; Lineberger Comprehensive Cancer Center, University of North Carolina at Chapel Hill, Chapel Hill, NC, USA

**Author notes:** Corresponding author: Department of Biology, University of North Carolina at Chapel Hill, 616 Fordham Hall, Campus Box 3280, Chapel Hill, NC 27599-3280.

**Keywords:** Tardigrades, BMP, FGF, Segmentation, Mesoderm

## Abstract

**Background:** A small number of signaling pathways regulates development in most animals, yet we do not know where these pathways are deployed in embryos of many animal phyla. Filling such gaps can contribute to understanding how diverse body shapes arise from differential deployment of conserved signaling pathways. Here, we examined where conserved pathways are deployed in tardigrades, a panarthropod phylum with miniaturized, segmented bodies.

**Results:** We used *in situ* mRNA detection in the tardigrade *Hypsibius exemplaris* to reveal expression patterns of FGF and BMP signaling pathway components during body segmentation and early mesoderm development. Among the patterns examined, we found an FGF ligand and receptor expressed near each other in segmentally iterated regions of ectoderm and endomesoderm, respectively. We also found a BMP ligand and antagonist expressed in dorsoventrally-restricted patterns in the lateral ectoderm.

**Conclusions:** The detected patterns suggested specific hypotheses for further research: possible FGF signaling between ectoderm and endomesoderm, and possible roles of BMP signaling in dorsal-ventral patterning of lateral ectoderm. We compared our results with published expression patterns for FGF and BMP pathways across panarthropods, to contribute to previous hypotheses for how the development of segments and mesoderm may have evolved in the emergence of this clade of diversly-shaped animals.

**Bullet Points:**

- Double mRNA detection in tardigrades revealed expression patterns of FGF and BMP pathway genes
- *fgf8* was detected primarily in ectoderm but absent from internal germ layers, and
- *fgfrl1* was detected in all layers and enriched in internal layers
- *dpp* was detected more dorsally than *sog* in lateral ectoderm
- *fgf8* and *dpp* gene expression exhibit different segmentally iterated domains in ectoderm
- Results suggest specific hypotheses for roles for FGF signaling and BMP signaling in tardigrade development

## Introduction

Embryonic development relies on precise temporal and spatial patterning to form specific body shapes. Organisms come in many unique shapes, as the result of unique developmental trajectories, yet each trajectory relies on widely conserved cell signaling pathways. Among animals, these include the FGF, BMP, EGF, TGF-β, Wnt, Hedgehog, Hippo, and Notch signaling pathways (Pires-daSilva and Sommer, 2003). There is a breadth of knowledge about how cell signaling refines developmental patterns in common model organisms (Pires-daSilva and Sommer, 2003). Much less is understood about how cell signaling facilitates the formation of other body forms. Such gaps limit our understanding of what is possible in the toolkit to build an embryo.

Addressing such gaps is possible with comparisons across developmental programs, particularly among closely related animals with differently shaped embryonic body plans. Ecdysozoa is a group of diversly shaped animals that includes the well-studied model systems *C. elegans* and *D. melanogaster*, as well as many emerging model systems (Giribet and Edgecombe, 2017; Telford et al., 2008). Panarthropoda is a subset of Ecdysozoa, those with segmented bodies, consisting of the arthropods (including *D. melanogaster*), onychophorans, and tardigrades (Howard et al., 2022; Telford et al., 2008). The repeating body segments of panarthropod animals are defined by a mostly homologous order of Hox gene expression in embryonic stages, yet panarthropods exhibit widely diverse adult structures (Akam, 1995; Franke and Mayer, 2014; Hughes and Kaufman, 2002; Janssen, 2017; Janssen et al., 2014; Peel et al., 2005; Smith et al., 2023, 2016; Smith and Goldstein, 2017). Such embryonic homology affords a powerful platform for building hypotheses of how different developmental trajectories transform these seemingly homologous segments into very different shapes. Positional information from cell signaling contributes to regulating differentiation of the germ layers and specifying the location, size, and morphology of tissue folds, organs, and appendages during development. Therefore, we can learn more about how homologous body segments are refined into a variety of structures by detecting where conserved pathways are deployed in body segments of species from across panarthropoda. Tardigrades comprise an entire phylum of panarthropod animals whose development is little studied compared to its arthropod relatives. We sought to uncover where conserved signaling pathways are deployed in a tardigrade species during and immediately following the emergence of body segments in embryos, to inform comparisons made across panarthropods.

The tardigrade body plan consists of a head with two eyes and four leg-bearing trunk segments and is uniquely miniaturized among panarthropods, with tardigrade body regions likely homologous to just the head and very posterior of arthropod and onychophoran bodies (Gross et al., 2019; Schill, 2018; Smith et al., 2016; Smith and Goldstein, 2017). One species of tardigrade, *H. exemplaris*, is an emerging model system, with established methods for *in situ* hybridization to visualize gene expression patterns (Gabriel et al., 2007; Goldstein, 2018; Goldstein and Blaxter, 2002; Heikes et al., 2023; Smith, 2018). Harnessing the spatial and temporal information provided from *in situ* hybridization, previous studies have shed light on mechanisms that may explain how the tardigrade’s segmented body was miniaturized from the panarthropod ancestor, by comparing expression patterns between tardigrades, onychophorans, and arthropods. These proposed mechanisms include 1) loss of an entire body region and associated hox genes, 2) loss of an intermediate leg region and associated leg gap gene with broader expression of remaining leg gap genes, and 3) loss of several Wnt ligand genes and a lack of segmentally iterated expression of the remaining Wnt ligands (Chavarria et al., 2021; Game and Smith, 2020; Smith et al., 2023, 2016). We took an approach also relying on *in situ* hybridization to reveal where members of conserved signaling pathways are deployed during early stages of endomesodermal and ectodermal segmentation, which reflect the first anatomical evidence of body segments in *H. exemplaris* embryos.

The Fibroblast Growth Factor (FGF) and Bone Morphogenetic Protein (BMP) conserved pathways are frequently deployed in refinement of endomesoderm and ectoderm across animal embryos, including members of Panarthropoda (Andrikou and Hejnol, 2021; Beermann and Schröder, 2008; Donoughe et al., 2014; Ewen-Campen et al., 2013; Fan et al., 2018; Green et al., 2013; Hamaratoglu et al., 2014; Hogvall et al., 2018; Huang and Stern, 2005; Janssen et al., 2015; Kainz, 2009; Kimelman and Kirschner, 1987; Matus et al., 2007; Muha and Müller, 2013; Nakamura and Extavour, 2016; Pechmann et al., 2017; Sharma et al., 2015; Stathopoulos et al., 2004; Suzuki et al., 1994; Treffkorn and Mayer, 2013; Van Der Zee et al., 2006; Wijesena et al., 2017; Winnier et al., 1995). BMP and FGF signaling also integrate with and feed back on one another during mesoderm development in animals, including panarthropods studied thus far (Kimelman and Kirschner, 1987; Luo, 2017; Row et al., 2018; Stathopoulos et al., 2004; Wang et al., 2023). Therefore, examining expression patterns of genes from both pathways may shed light on spatiotemporal refinement of ectoderm and mesoderm in the body segments of *H. exemplaris* embryos.

Using double fluorescence *in situ* hybridization (FISH) to simultaneously detect expression patterns of two genes at a time in embryos of the tardigrade *H. exemplaris*, we revealed the spatiotemporal expression of FGF and BMP pathway genes during the processes of endomesodermal segmentation and subsequent ectodermal segmentation. Based on these observed patterns and known roles in other animals, we present hypotheses about where genes may have been expressed in the ancestral panarthropods and about the potential roles of FGF and BMP signaling in refining tardigrade body shape during segmentation and mesoderm development, which will be testable upon expansion of genetic tools for use in tardigrade embryos.

## Results and Discussion

### Identification of conserved signaling pathway genes in *H. exemplaris*

We identified putative homologs for genes encoding members of the FGF and BMP signaling pathways in *H. exemplaris*. We searched for upstream pathway components and putative target genes, as these are most likely to be differentially expressed. We identified all likely homologs in published *H. exemplaris* transcriptomes (Levin et al., 2016; Yoshida et al., 2017) by NCBI’s tBLASTn search, using *D. melanogaster* protein sequences as queries (Kent et al., 2002) and validating by reciprocal BLASTp against the *D. melanogaster* protein database to confirm that identified homologs returned the original query (Altschul et al., 1990; Gerts et al., 2006; Sayers et al., 2022). We analyzed top hits further with protein domain database tools to confirm protein structures matched those of the queries (Letunic et al., 2021; Letunic and Bork, 2018). We sought further support of gene homology for top hits of query protein sequences through maximum likelihood phylogenetic reconstruction of likely sequence evolution among homologs and orthologs from other ecdysozoan and non-ecdysozoan species, with branch support from 500 bootstraps [details in Experimental Procedures] (Kumar et al., 2018). Support varied from gene to gene, and we tentatively assigned gene names based on the support we found and the BLAST results, domain analyses, and alignments; below each phylogenetic tree, we show an alignment of the sequences used to build the trees, capturing a subset of the species represented in each tree [Supp. Figs. 2-9]. We consulted a published staged transcriptome through *H. exemplaris* development to aid in prioritizing genes for mRNA detection by FISH at our developmental stages of interest – endomesodermal and ectodermal segmentation (Levin et al., 2016).

For the FGF pathway in published *H. exemplaris* transcriptomes, we identified one FGF ligand homolog: *fgf8* (Supp. Fig. 1) and two homologs of FGF receptors: *fgfrl1* and *fgfrl2* (FGF receptor-like 1 and 2, respectively, Supp. Fig. 2). Expression levels in the published staged transcriptome suggested to us that *fgfrl1* is expressed at high enough levels for detection by fluorescence *in situ* hybridization (FISH), but *fgfrl2* is very weakly expressed (Levin et al., 2016). Our preliminary staining by chromogenic *in situ* hybridization also aligned with this prediction, as we detected low to no signal of *fgfrl2* during segmentation stages (data not shown). We concluded that *fgfrl1* is likely to be relevant during these stages of embryonic development in *H. exemplaris* and moved forward with this gene to identify cells likely to be receptive to FGF signaling. The *H. exemplaris* homolog of *fgf8* was not represented in the staged transcriptome, so we could not consult it for expected expression levels. We also identified one homolog of the mesodermal transcription factor *snail* in hopes of using it as a marker of embryonic mesoderm and potential FGF responsive cells, since FGF is known to regulate mesoderm development (Supp. Fig. 3). This *snail* homolog was weakly expressed in the staged transcriptome (Levin et al., 2016).

For the BMP pathway in published *H. exemplaris* transcriptomes, we found one homolog each of the BMP2/4-type ligand *decapentaplegic* (*dpp*), the BMP5/6/7/8-type ligand *glass bottom boat* (*gbb*) (Supp. Fig. 4), the extracellular BMP antagonist *short gastrulation* (*sog*) (Supp. Fig. 5), the Type 1 and Type 2 BMP receptors (Supp. Fig. 6) *thickveins* and *punt*, the Type 1 BMP receptors *saxophone* and *baboon*, and the Sog-cleaving protease *tolloid* (Supp. Fig. 7). We also identified one homolog each of known downstream transcriptional targets of BMP signaling in arthropods as potential markers of BMP responsive cells: *dorsocross1* (*doc1*) and *eyes absent* (*eya*) (Supp. Fig. 8 and Supp. Fig. 9) (Dominguez et al., 2016). FGF signaling has also been found to target *eya* expression (Ahrens and Schlosser, 2005). All of these identified homologs were highly expressed between elongation and segmentation stages in the published staged transcriptome (Levin et al., 2016). Another known target of BMP signaling in arthropods, *c15* (*clawless*), was cloned in a previous publication, which reported its expression domains at post-segmentation stages in developing limbs (Mapalo et al., 2024).

We refer to these genes by their likely homologs’ names moving forward. Using FISH, we detected where each of these upstream signaling pathway genes and putative target genes is expressed during segmentation of *H. exemplaris* embryos. Although mRNA distributions are affected by mRNA synthesis, localization, persistence, and depletion, we refer to the presence of mRNA, detected by FISH, as gene expression for simplicity. A summary of the relevant embryonic stages examined is presented in Fig. 1. In the sections that follow, we present detected expression of components of FGF signaling first and of BMP signaling second.

**Figure 1.**
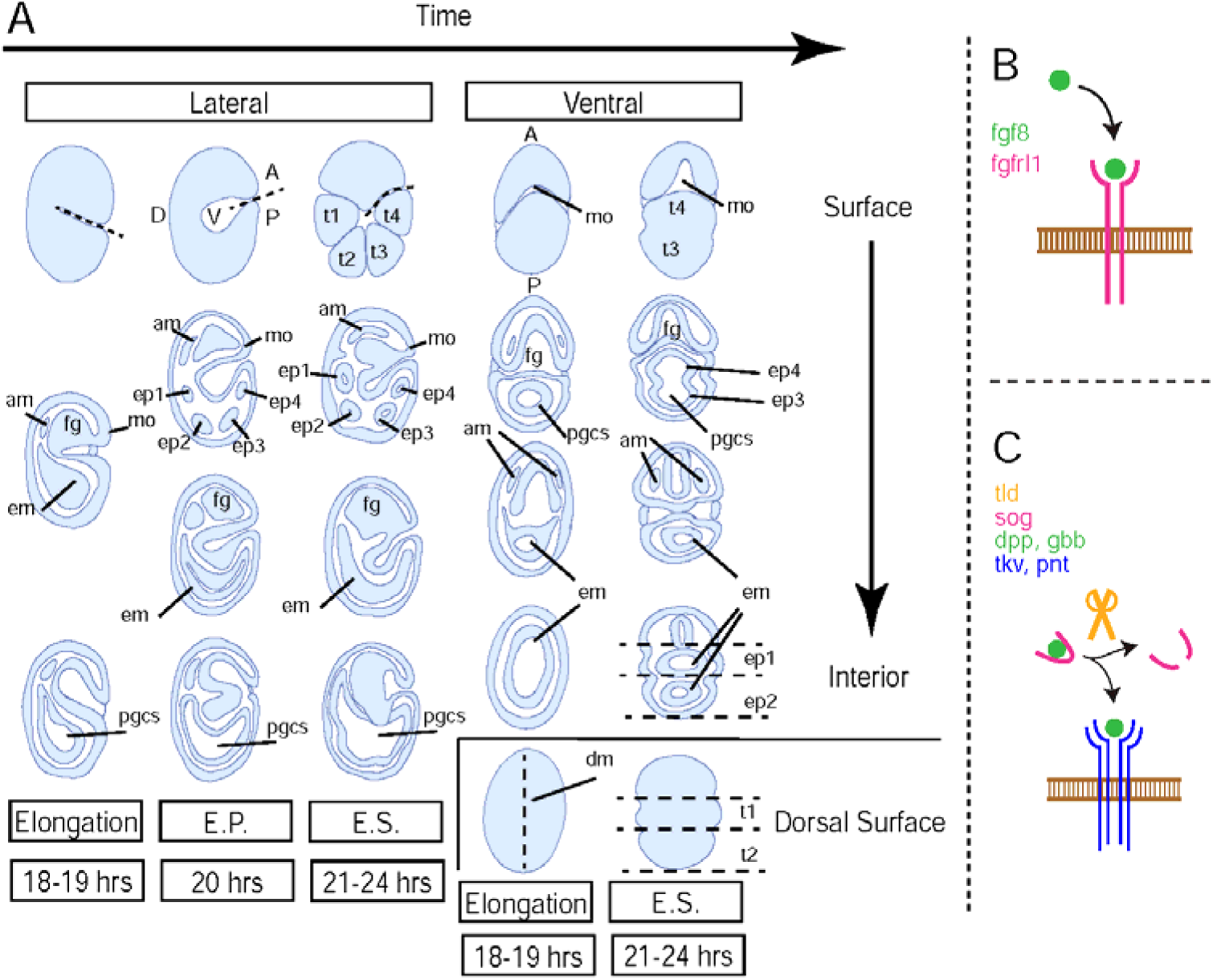
Introduction to morphology of *Hypsibius exemplaris* embryos between elongation and segmentation stages. A. Drawings representative of *H. exemplaris* embryos at elongation, endomesodermal pouch formation (E.P.), and ectodermal segmentation (E.S.) stages, shown from right lateral and ventral views. From top to bottom, images show surface to internal views (and dorsal surface for ventral views). Anterior (A) is up and Posterior (P) is down in all drawings. Dorsal (D) and (V) are to the left and right, respectively, in the lateral views. Certain morphological features are labelled: mouth (mo), anterior mesoderm (am), dorsal midline (dm), foregut (fg), endomesoderm (em), endomesodermal pouch (ep), trunk segment (t), and primordial germ cells (pgcs). B. Schematic of FGF signaling showing extracellular ligand (green) binding transmembrane receptors (magenta), which dimerize. C. Schematic of BMP signaling showing extracellular ligand (green) binding extracellular antagonist (magenta), which is cleaved by extracellular protease (orange), free the ligand to bind transmembrane receptors (blue), which form a heterotetramer.

### By ectodermal segmentation, FGF ligand *fgf8* is expressed in segmentally iterated patches of ectodermal cells and FGF receptor *fgfrl1* is enriched in underlying endomesodermal pouches

To reveal likely FGF-sending and FGF-receiving cells, we simultaneously detected where *fgf8* and *fgfrl1* are expressed at 24 hours post laying (hpl) when ectodermal and endomesodermal segments are both anatomically apparent. Among arthropods studied thus far, gene homologs of the FGF ligand *fgf8* are expressed in or near the developing foregut, in a region of the head segment that defines the midbrain-hindbrain boundary, and in a posterior growth zone in species that undergo posterior growth, which then persists in the posterior of body segments as they form from the growth zone (Beermann and Schröder, 2008; Kainz, 2009; Muha and Müller, 2013; Stepanik et al., 2020). The fruit fly *D. melanogaster* does not undergo posterior growth but still exhibits segmental expression of its *fgf8* homologs in ectoderm (Stepanik et al., 2020). Current evidence suggests that onychophorans do not undergo posterior growth, and FGF signaling has yet to be studied in an onychophoran system (Hogvall et al., 2014; Mayer et al., 2010). Tardigrades also do not undergo posterior growth (Hejnol and Schnabel, 2005; Smith et al., 2016; Smith and Goldstein, 2017). We found that *fgf8* was expressed in a segmentally iterated pattern in the ectoderm – in patches of lateral ectodermal cells in the posterior of each trunk segment on both the left and right sides of embryos (Fig. 2 and Video 1). A patch of *fgf8* expression was detected laterally near the middle of the head segment in ectoderm, potentially contributing to regional patterning of the brain, which may still occur at the level of cell fates within the unipartite brain structure of tardigrades (Smith et al., 2018). Another patch of *fgf8* expression was detected in a small set of cells at the posterior of the head segment in ectoderm and endomesoderm, likely where the esophagus develops from foregut (Fig. 2). While *fgf8* was expressed in a segmentally repeating pattern in the ectoderm, *fgfrl1* was expressed broadly in the embryo and enriched in the endomesodermal pouches found within each of the four trunk segments, in the developing foregut in the head segment, and to a lesser extent in the ectoderm (Fig. 2 and Video 2). This is consistent with a possibility that all mesoderm is receptive to FGF8 signal, and that FGF8 might pattern the underlying *fgfrl1*-expressing mesoderm in addition to *fgfrl1*-expressing ectoderm. Segmentally iterated posterior patches of *fgf8* may be ancestral to panarthropod species; however, addressing this demands data from a larger sampling of panarthropods, especially onychophorans.

**Figure 2.**
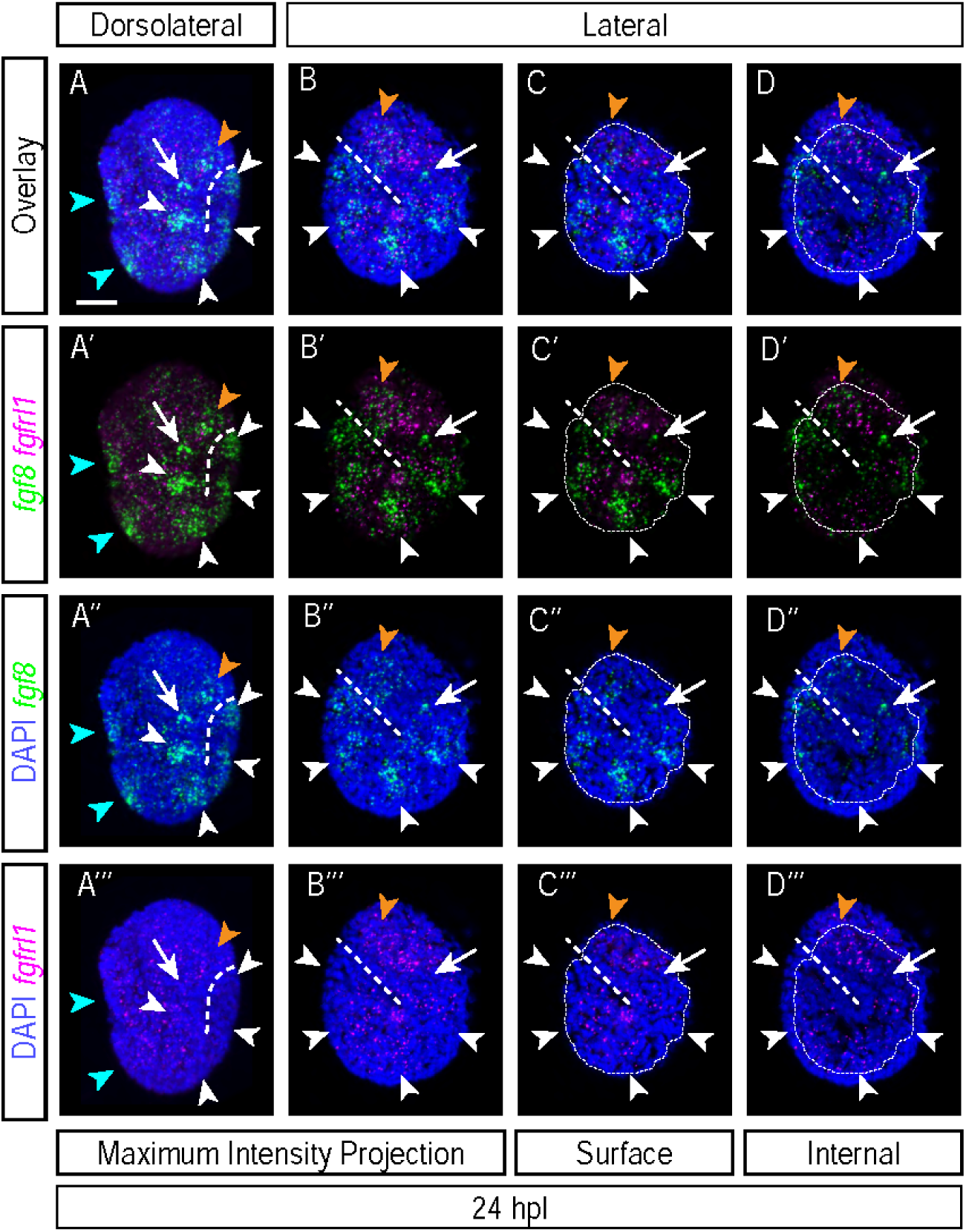
Expression patterns of *fgf8* and *fgfrl1* mRNAs at 24 hpl reveal ectodermal patches of *fgf8* in segment posteriors and global expression and endomesodermal enrichment of *fgfrl1*. A-D. Expression patterns of *fgf8* and *fgfrl1* at 24 hpl (n=5 experiments, 8, 3, 4, 2, and 2 embryos respectively). A-B. Maximum intensity projections of embryos from a dorsolateral (A) and lateral view. C. Projection of uppermost layers of the embryo in B. D. Projection of internal layers of the embryo in B. Small region outlined with a dashed line indicates the upper layers projected in C, which are omitted in the internal layers projected in D. Arrowheads indicate ectodermal patches of *fgf8* signal in trunk segment posteriors. Blue arrowheads indicate two of the ectodermal patches of *fgf8* signal running along the left side of the embryo in A. Orange arrowheads indicate the anterior head segment patch of *fgf8* signal. Arrows indicate small groups of ectodermal and endomesodermal cells enriched for *fgf8* at the posterior of head segments. Right dorsolateral view in A. Left lateral view in B-D. Scale bar = 10 μm.

We next wondered how early these patches of *fgf8* emerge relative to the anatomical appearance of body segments, since by 24 hpl, body segments are 3 hours old (Fig. 1 A).

### Segmentally iterated *fgf8* expression appears prior to ectodermal segmentation and near the time when endomesodermal pouches form

To determine if segmentally iterated *fgf8* expression preceded, arose concomitantly with, or followed body segmentation, we detected expression at earlier stages – at late elongation (∼18 and 19 hpl) before segments are apparent, at endomesodermal pouch formation (∼20 hpl) when endomesoderm begins to appear segmentally divided, and at early ectodermal segmentation (∼21-22 hpl) when ectoderm begins to appear segmentally divided (Fig. 1 A). These stages correspond to stages 11-13 from previous publications (Chavarria et al., 2021; Gabriel et al., 2007). Between 18 hpl and 22 hpl, *fgfrl1* was enriched in endomesodermal cells, and this pattern did not change dramatically. In contrast, the number of ectodermal domains expressing *fgf8* increased over time in this period. At elongation stages, there were fewer, broad patches of ectodermal *fgf8* expression than at later stages (Fig. 3 A-A’’’ and Supp. Fig. 10). Additionally, the patches of cells expressing *fgf8* appeared first in the anterior half of the embryo at elongation stages and only later when endomesoderm was segmented did they line the embryo from anterior to posterior. Following or concomitant with ectodermal segmentation (∼21-22 hpl), ectodermal patches of *fgf8* expression began to resemble what we saw at 24 hpl, i.e. segmentally iterated, in lateral pairs in the posterior of each trunk segment and in the middle of the head segment, in addition to expression in a small group of cells in the posterior of the head segment (Fig. 3 C-C’’’ and D-D’’’). Ectodermal segmentation is evident by the presence of furrows in the ectoderm at the approximate boundaries between body segments (Fig. 1) (Chavarria et al., 2021; Gabriel, 2007; Gabriel et al., 2007; Gabriel and Goldstein, 2007). At 20 hpl, when endomesodermal pouches have formed but ectodermal segmentation has not occurred, patches of *fgf8* expression were present in the lateral ectoderm on each side of the embryo (Fig. 3 B-B’’’). This expression pattern reveals that enrichment of *fgf8* in the ectoderm of each segment precedes apparent anatomical segmentation of ectoderm and either follows or occurs concomitantly with segmentation of endomesoderm.

**Figure 3.**
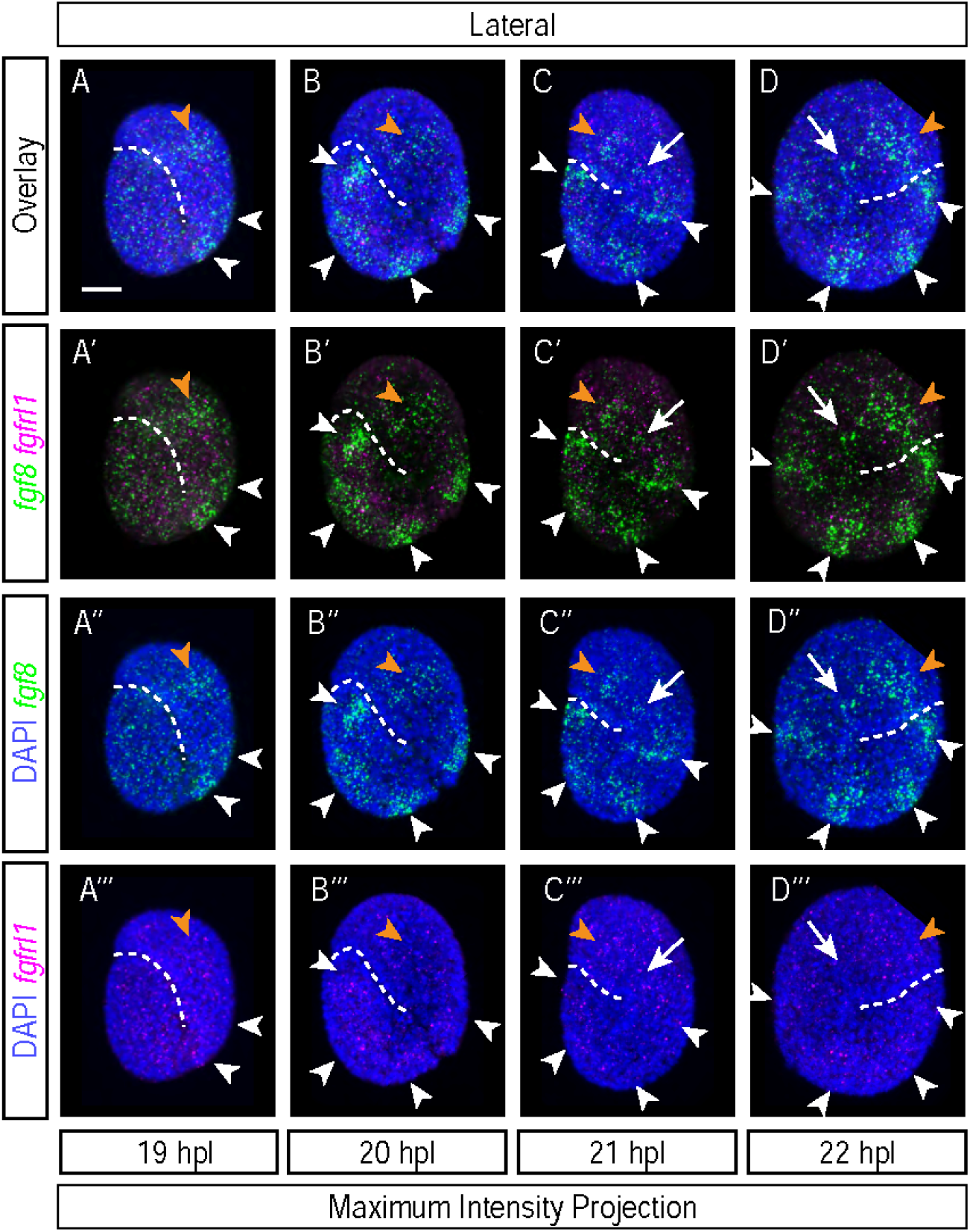
Lateral ectoderm patches of *fgf8* mRNA arise prior to ectodermal segmentation and near the time of endomesodermal pouch formation. A-D. Expression patterns of *fgf8* and *fgfrl1* mRNAs at 19 hpl (A, n=2 experiments, 8 and 4 embryos) 20 hpl (B, n=3 experiments, 9, 8, and 6 embryos) 21 hpl (C, n=5 experiments, 8, 11, 6, 5, and 7 embryos) and 22 hpl (D, n=5 experiments, 8, 3, 4, 2, and 2 embryos). All images are maximum intensity projections of embryo from lateral views. Arrowheads indicate ectodermal patches of *fgf8* signal. Orange arrowheads indicate the patch of *fgf8* signal in the anterior of head segments. Arrows indicate small groups of cells enriched for *fgf8* at the posterior of head segments. Left lateral view in A-C. Right lateral view in D. Scale bar = 10 μm.

FGF plays a role in posterior segment polarity in deuterostomes, along with Wnt. Thus far, evidence in ecdysozoans, a protostome superclade, only suggests a role of Wnt in posterior segment polarity, not FGF (Angelini and Kaufman, 2005; McGregor et al., 2008; Miyawaki et al., 2004). FGF does have a functional role in segment number in the beetle *Tribolium castaneum* and in segment size in the cricket *Gryllus bimaculatus*, but these roles are difficult to untangle from FGF’s function in body extension in these arthropods that undergo posterior growth (Kainz, 2009; Sharma et al., 2013). Expression of *fgf8* in segment posteriors across arthropods and in a tardigrade suggest this may be an unidentified role for FGF in panarthropods, but ruling this out confidently demands data from a larger representation of panarthropod species. If FGF8 is a segment polarity cue in tardigrades, then endomesodermal pouch formation might follow *fgf8* mRNA patch formation. We were unable to resolve the order in which these events occur, as we never saw an embryo with patches of *fgf8* mRNA but an unsegmented endomesoderm or vice versa. This would require a study using a live fluorescent reporter of *fgf8* mRNA, such as the MS2-MCP system, and DIC imaging of embryo morphology (Bertrand et al., 1998; Garcia et al., 2013; Lucas et al., 2013; Wu et al., 2012). Related to this, another study found by antibody staining that Engrailed protein, which is also expressed in segment posteriors, appears only after formation of endomesodermal pouches (Gabriel and Goldstein, 2007). Therefore, FGF8 might be an instructive cue for Engrailed expression, further reinforcing the posterior boundary of segments.

The patches of *fgf8* expression also appeared to change in orientation relative to the axes of the embryo. Upon ectodermal segmentation, the patches were positioned in cells in the posterior of each segment and had elongated forms that were aligned perpendicularly to the anterior-posterior axis (Fig. 2 and Fig. 3 C and D). Prior to ectodermal segmentation, the patches were elongated parallel to the anterior-posterior axis, along the segments (Fig. 3 B). Therefore, the pattern of *fgf8* is dynamic, or the *fgf8-*expressing cells move or change shape, during ectodermal segmentation of the embryo. FGF8 often serves as a guidance cue for mesodermal migration, and if this is the case in *H. exemplaris*, then this change in orientation of the *fgf8* patches may be an important directional cue (DeVore et al., 1995; Gryzik and Muller, 2004; Kadam et al., 2012; Klingseisen et al., 2009; Röttinger et al., 2008; Sharma et al., 2015). Alternatively or in addition, the underlying mesoderm, as it migrates, may feed back on *fgf8* to influence the direction of the patches and the formation of ectodermal segments, which appears to occur concomitantly with the re-orienting of *fgf8* patches, though determining this also requires the use of live imaging tools.

### The mesodermal transcription factor *snail* is expressed in non-mesodermal cells

To determine more precisely which cells in endomesodermal pouches contribute to the mesoderm, we detected expression for the ectodermal transcriptional repressor *snail*, which historically serves as a marker of mesodermal fate in animal embryos, together with the pro-mesodermal transcription factor *twist* (Leptin, 1991). However, *snail* was expressed throughout the embryo, in both ectodermal and endomesodermal layers of cells at both elongation and ectodermal segmentation stages (Fig. 4). Expression of *snail* appeared to become more enriched in the internal endomesodermal cells by 24 hpl but was still present in ectodermal cells (Fig. 4 A-D). Therefore, *snail* expression may not be a reliable marker of mesodermal fate at this stage in development. By double FISH, *snail* was expressed in cells that expressed *fgfrl1* (Fig. 4 A-A’’’ and B-B’’’) and in cells that expressed *fgf8* (Fig. 4 C-C’’’ and D-D’’’). Given this overlap, this expression pattern suggests that *snail* is expressed in both FGF sending and FGF receiving cells.

**Figure 4.**
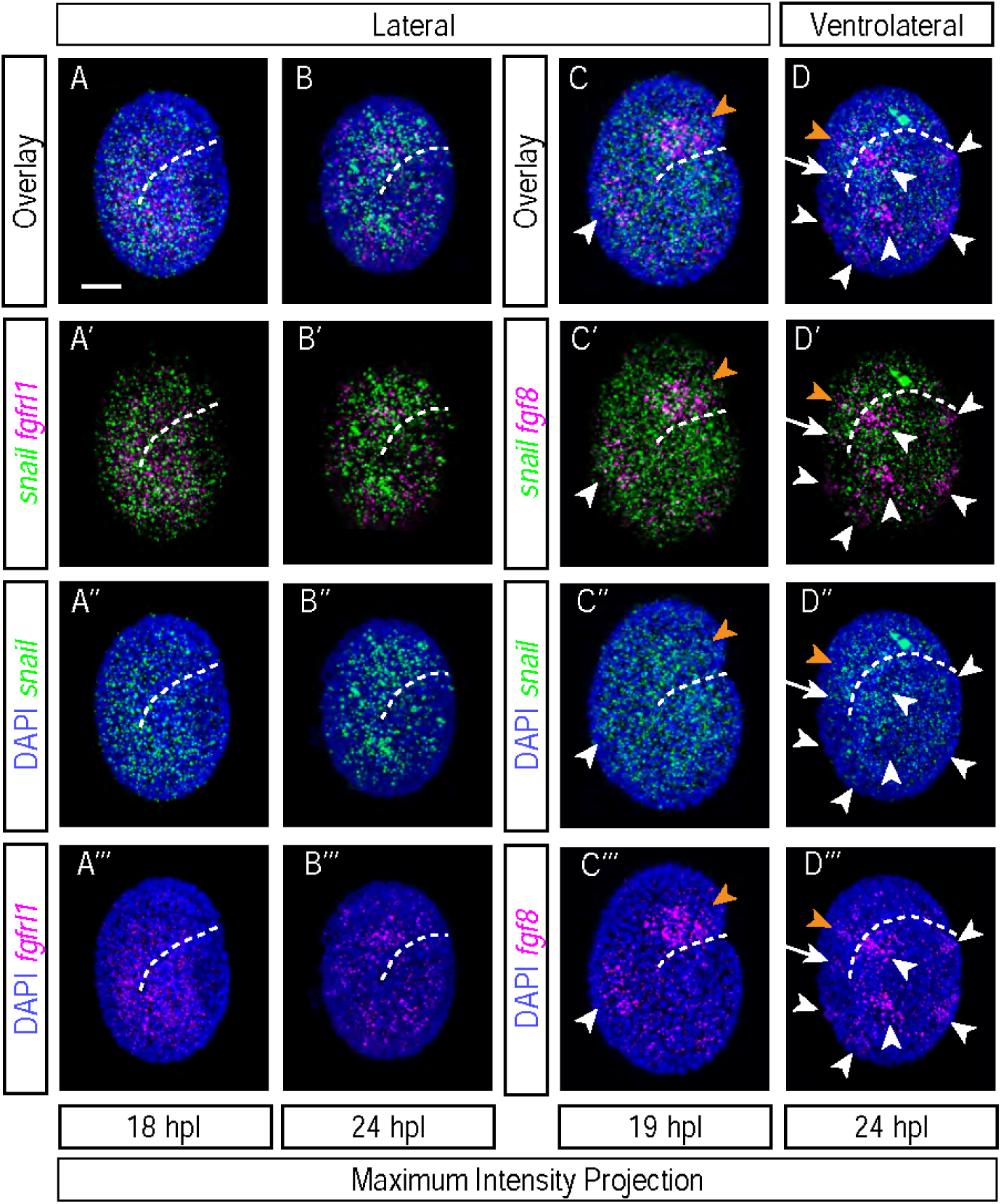
mRNA of the mesodermal transcription factor *snail* is present in non-mesodermal cells. A-B. Enrichment patterns of *snail* and *fgfrl1* mRNAs at 18 hpl (A, n=2 experiments, 8 and 2 embryos) and 24 hpl (B, n=2 experiments, 1 and 3 embryos). C-D. Enrichment patterns of *snail* and *fgf8* mRNAs at 19 hpl (C, n=2 experiments, 8 and 6 embryos) and 24 hpl (D, n=2 experiments, 8 and 5 embryos). All images are maximum intensity projections. Arrowheads indicate ectodermal patches of *fgf8* signal. Orange arrowheads indicate the patch of *fgf8* signal in the anterior of head segments. Arrows indicate small groups of cells enriched for *fgf8* at the posterior of head segments. Right lateral view in A-C. Right ventrolateral view in D. Scale bar = 10 μm.

This expression pattern of *snail* was surprising, given that in other systems, Snail serves as a marker and promoter of mesoderm fate through inhibition of non-mesodermal genes (Alberga et al., 1991; Cano et al., 2000; Chopra and Levine, 2009; Simpson, 1983). Although it does not appear to be a reliable marker of mesodermal fate at these stages in *H. exemplaris* development, the expression pattern of *snail* is potentially reflective of aspects of mesodermal development in this system. It is possible that either mesoderm migrates from many starting points in development, including the ectoderm layer, or that many nonmesodermal cells express *snail* and only later in development is *snail* restricted to mesoderm. It is also possible that *snail* instead marks neuroectoderm as it internalizes from the surface of the embryo, as *snail* marks neuroectoderm in many panarthropods and other animal systems studied (Erives et al., 1998; Hannibal et al., 2012; Hemavathy et al., 1997; York et al., 2019).

Based on all of our results examining expression of FGF pathway genes, we conclude that the ligand-encoding gene *fgf8* is expressed primarily in lateral ectoderm in the posterior of each body segment, as well as in the head, and the receptor-encoding gene *fgfrl1* is expressed broadly and enriched in endomesoderm, including in pouches underlying each ectodermal *fgf8*-expressing patch of cells.

### BMP ligand *dpp* and antagonist *sog* are expressed in lateral ectoderm, and *dpp* is expressed more dorsally than *sog*

To determine where components of the BMP signaling pathway are expressed at an ectodermal segmentation stage (24 hpl), we detected expression of BMP receptors *tkv* and *punt* (which frequently heterodimerize and bind *dpp*), BMP ligands *dpp* and *gbb*, BMP antagonist *sog*, and Sog-cleaving protease *tld* (Mullins, 1998; Nunes da Fonseca et al., 2010; Ramel and Hill, 2012). By single FISH, receptors *tkv* and *punt* were ubiquitously expressed across the embryo, consistent with a possibility that all cells may be receptive to BMP signaling (Supp. Fig. 11). However, the precise pattern of cells receptive to BMP signaling via Gbb and/or Dpp may be further refined by the expression patterns of other BMP receptor genes, such as *saxophone* and *baboon*. BMP5/7/8-type ligand Gbb regulates development of the germ cells in the arthropod *G. bimaculatus*, as well as in vertebrate systems, but so far BMP signaling has not been implicated in germ cell development in other panarthropods (Donoughe et al., 2014; Lawson et al., 1999; Lochab and Extavour, 2017). In *H. exemplaris*, the ligand *gbb* was ubiquitously expressed throughout most of the embryo (Supp. Fig. 12). However, *gbb* expression appeared to be absent from the cells previously defined as the primordial germ cells (Heikes et al., 2023) but present in somatic cells surrounding the primordial germ cells (Supp. Fig. 12), whereas receptors *tkv* and *punt* were expressed in the primordial germ cells (Supp. Fig. 11). This expression pattern is consistent with a possibility that signaling by Gbb might regulate primordial germ cell development in this tardigrade species.

In arthropods and onychophorans, BMP2/4-type ligand gene *dpp* is expressed along the trunk near trunk ganglia, near the developing mouth, and in the anlage of limbs, and is known to regulate embryo dorsal-ventral patterning, mesoderm differentiation, nervous system development, and appendage patterning (Donoughe et al., 2014; Morisato and Anderson, 1995; Nakayama et al., 2000; Ramel and Hill, 2012; Winnier et al., 1995). By double FISH, the ligand *dpp* and the antagonist *sog* were expressed in ventro-lateral ectoderm (Fig. 5 and Video 3). Neither gene was appreciably expressed in the most dorsal regions of the embryo (Fig. 5 C and Video 4). *dpp* was expressed in more dorsal regions of lateral surfaces than *sog* (Fig. 5 C). This lateral enrichment of *dpp* and *sog* is consistent with a possible role for Dpp in dorsal-ventral patterning of lateral ectoderm, perhaps by elaborating fates along the dorsal-ventral axis, rather than symmetry breaking, as symmetry breaking in the dorsal-ventral axis has already occurred by this stage of *H. exemplaris* development. Alternatively or in addition, this expression pattern is consistent with a potential role for Dpp in regulating development of the neural trunk ganglia that run in parallel rows from anterior to posterior along the animal’s ventral surface.

**Figure 5.**
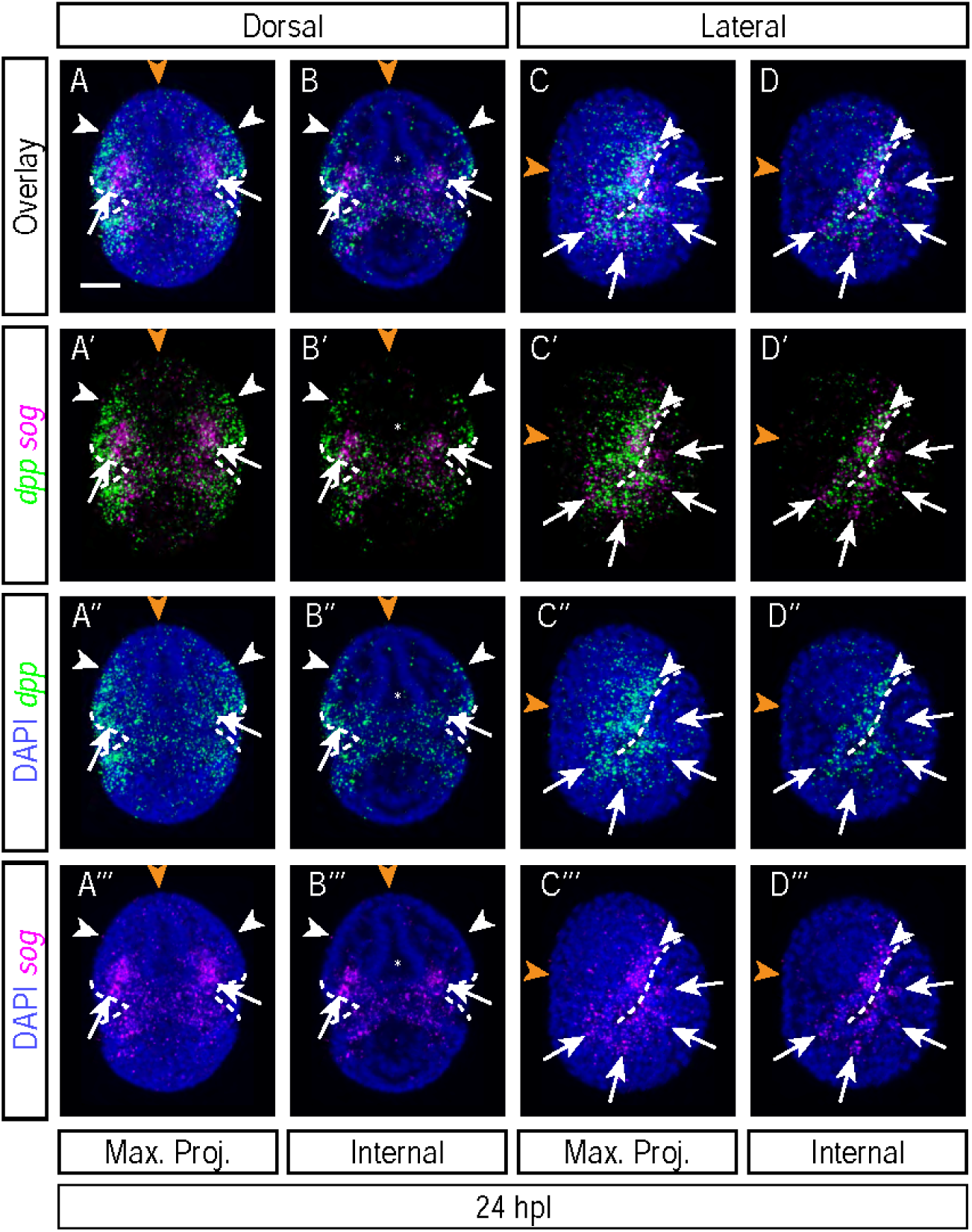
mRNAs of BMP ligand *dpp* and antagonist *sog* are expressed in lateral ectoderm, with *dpp* expressed more dorsally than *sog.* A-D. Enrichment patterns of *dpp* and *sog* mRNAs at 24 hpl (n=6 experiments, 6, 1, 1, 1, 2, and 13 embryos). A and C Maximum intensity projections of embryos from a dorsal (A) and lateral view. B. Projection of internal layers of the embryo in A. D. Projection of internal layers of the embryo in C. Arrowheads indicate the extent of spread of *dpp* mRNA dorsally. Orange arrowheads indicate the absence of enrichment for *dpp* and *sog* at the dorsal midline of the embryo. Arrows in A-B indicate bands of cells enriched for *sog* on either side of the developing mouth. Arrows in C-D indicate the endomesodermal cells enriched for *sog* in each endomesodermal pouch. Asterisks in A-B indicate the hollow space within the developing foregut, which is not enriched for *sog* in its epithelium. Dorsal view in A and B. Right lateral view in C and D. Scale bar = 10 μm.

At the anterior-most tip of the head where the mouth likely develops, *sog* was expressed in bands on either side laterally but did not appear to be expressed in the epithelium of the developing foregut (Marcus, 1929) (Fig. 5 A and B). *dpp* was expressed in large patches of ectodermal cells overlapping and dorsally adjacent to these two bands of cells enriched for *sog* expression (Fig. 5 A and B). The expression of *sog* on either side of the developing mouth led us to speculate that Dpp might provide signals that promote or inhibit mouth formation, akin to Dpp’s role in regulating formation of appendages among arthropods studied. *sog* was also expressed in one endomesodermal cell on the ventral side of each of the endomesodermal pouches in each body segment (Fig. 5 D). *dpp* was expressed in segmentally iterated bands extending over each segment near the segment boundaries (Fig. 5 C and D). These bands were in the overlying ectoderm adjacent to the endomesodermal cells that expressed *sog*. Due to the banded expression pattern of *dpp* along each segment, we hypothesize that Dpp might regulate the dorsal-ventral position of future limbs in each trunk segment, similar to the role of Dpp in regulating appendage formation in arthropods and onychophorans (Goto and Hayashi, 1997; Janssen et al., 2015; Spencer et al., 1982). Alternatively or in addition, we hypothesize that Dpp might regulate cuticle patterning by guiding cuticle deposition or sites of cuticle furrowing. Finally, due to the role of ectodermal BMP in regulating mesoderm development in *D. melanogaster*, *Danio rerio*, and *M. musculus*, among other animals, and the ectodermal enrichment of *dpp*, we hypothesize that Dpp might regulate mesoderm development from overlying ectoderm (Frasch, 1995; Row et al., 2018; Staehling-Hampton et al., 1994).

We wondered whether *dpp* and *sog* expression in these regions preceded or coincided with the segmentation of the ectoderm and/or endomesoderm. They could also correspond with the future positions of the limb buds or with developing ventral trunk ganglia, both known functions of DPP signaling in other panarthropods studied (Holley et al., 1995; Janssen et al., 2015; McKay et al., 2009; Minelli et al., 2013; Spencer et al., 1982; Stollewerk, 2016; Treffkorn and Mayer, 2013; Vargas-Vila et al., 2010).

We compared expression of *dpp* and *sog* at earlier stages: elongation (18 and 19 hpl), endomesodermal pouch formation (20 hpl), and ectodermal segmentation (21 hpl) (Fig. 1 A). Lateral expression of *dpp* and *sog* were apparent by elongation (18 hpl) (Fig. 6 A). However, *dpp* was expressed in broader swaths of lateral ectoderm at elongation than at segmentation stages, and *sog* was expressed in a shorter region along the A-P axis of lateral ectoderm at elongation than at segmentation stages (Fig. 6 A-B vs C-D). Bands of *dpp* expression either coincided with or followed ectodermal segmentation (Fig. 6 D), and endomesodermal cells began expressing *sog* after endomesodermal pouch formation and during or after ectodermal segmentation (Fig. 6 C-D). Antero-lateral bands of *sog* and *dpp* near the developing mouth were apparent by elongation stages (18 hpl) (Fig. 6 A-B). These patterns indicate that *dpp* and *sog* are already expressed laterally prior to the processes of endomesodermal pouch formation and ectodermal segmentation and that the patterns of *dpp* and *sog* expression change throughout these morphological transformations.

**Figure 6.**
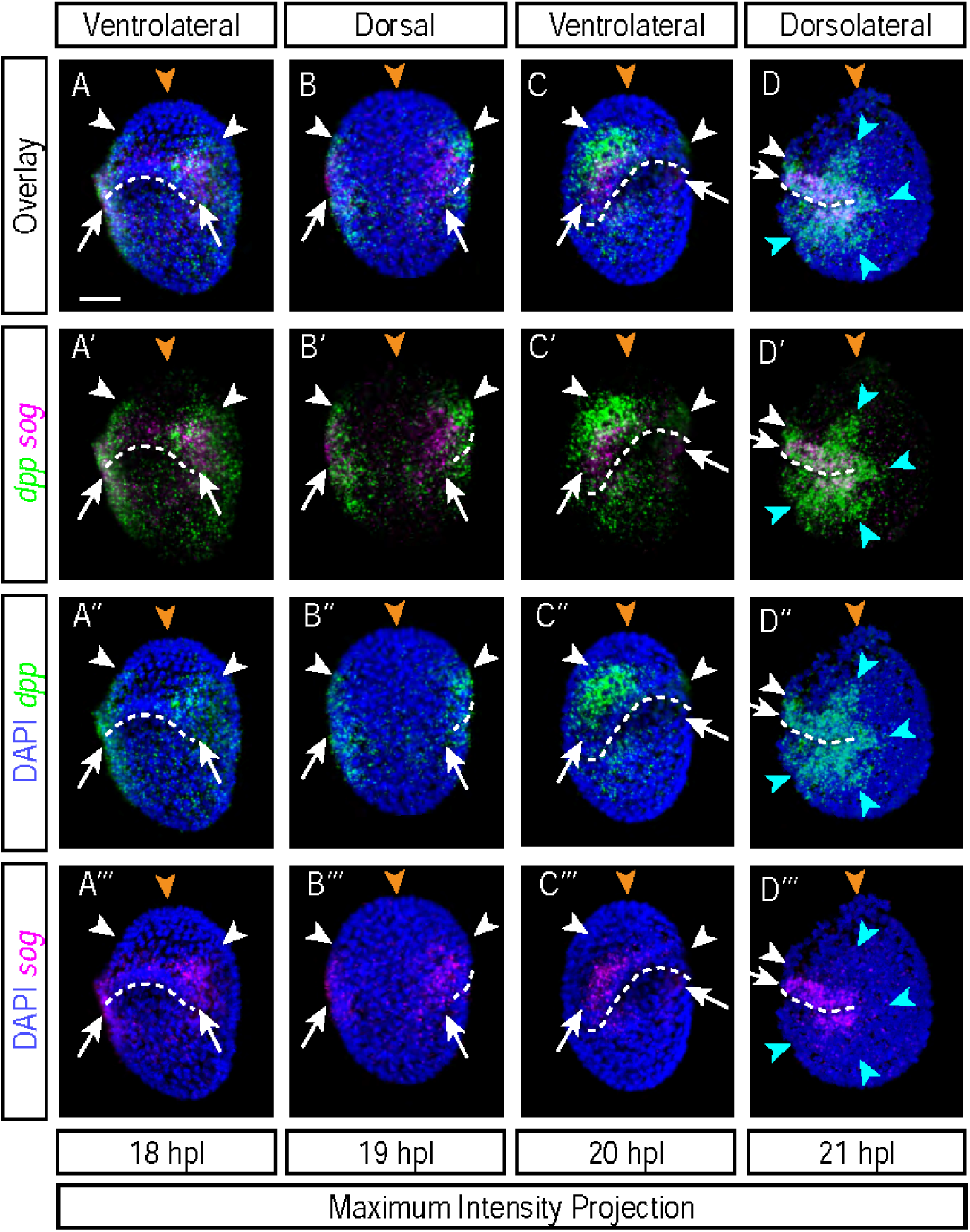
Expression of *dpp* and *sog* mRNAs changes between elongation and segmentation. A-D. Enrichment patterns of *dpp* and *sog* mRNAs at 18 hpl (A, n=3 experiments, 13, 2, and 1 embryos) 19 hpl (B, n=1 experiment, 7 embryos) 20 hpl (C, n=1 experiment, 7 embryos), and 21 hpl (D, n=3 experiments, 10, 16, and 2 embryos). All images are maximum intensity projections. Arrowheads indicate the extent *dpp* mRNA expression dorsally. Orange arrowheads indicate the absence of enrichment for *dpp* and *sog* at the dorsal midline. Arrows indicate bands of cells enriched for *sog* on either side of the developing mouth. Blue arrowheads indicate bands of *dpp* running over each segment. Left ventrolateral view in A. Dorsal view in B. Right ventrolateral view in C. Left dorsolateral view in D. Scale bar = 10 μm.

Sog might act to restrict spread of Dpp ligand in one or multiple axes. This is complicated by the varying impacts that Sog/Chordin can have on BMP signaling and spreading (Nunes da Fonseca et al., 2010; Shimmi et al., 2005); it is possible that Sog acts to facilitate further and directed spread of Dpp protein, in conjunction with Tolloid (Tld) and Twisted Gastrulation (Tsg) (Nunes da Fonseca et al., 2010; Shimmi and O’Connor, 2003; Van Der Zee et al., 2006).

### Expression of the protease *tld* is increasingly restricted along the anterior-most end of the head from 19 to 24 hpl

Having found *dpp* expression in lateral ectoderm and in bands near segment boundaries, and antagonist *sog* expression in a more ventrally restricted domain of lateral ectoderm than *dpp*, we wondered where mRNA of the Sog protein-cleaving protease Tld was expressed, as this impacts spread of Dpp extracellularly (Nunes da Fonseca et al., 2010; Shimmi and O’Connor, 2003; Van Der Zee et al., 2006). At elongation (19 hpl), *tld* was expressed in a wide band of cells along the anterior-most end of the head (Fig. 7 A). This band was mostly non-overlapping with the bands of *sog* expression that run on either side of the developing mouth (Fig. 5, Fig. 6, and Fig. 7 A) By endomesodermal pouch formation (20 hpl), the band of cells that expressed *tld* was thinner than at 19 hpl, indicating that *tld* expression becomes more restricted and/or that the *tld*-expressing cells condense in space between these stages (Fig. 7 B). This band of *tld* expression persisted through ectodermal segmentation (24 hpl) and remained mostly non-overlapping with the two bands of *sog* expression on either side (Fig. 7 C). At ectodermal segmentation stage, *tld* expression was mostly non-overlapping with the antero-lateral patches of cells that expressed *dpp*, which were located adjacent to and slightly overlapping with the bands of *sog* expression (Fig. 7 D and Fig. 5 and Fig. 6). *tld* did not appear to be expressed in the epithelium of the developing foregut (Fig. 7 C and D). These patterns indicate that *tld* is expressed in the cells where the mouth likely develops, surrounded on either side by cells that express *sog* and *dpp*.

**Figure 7.**
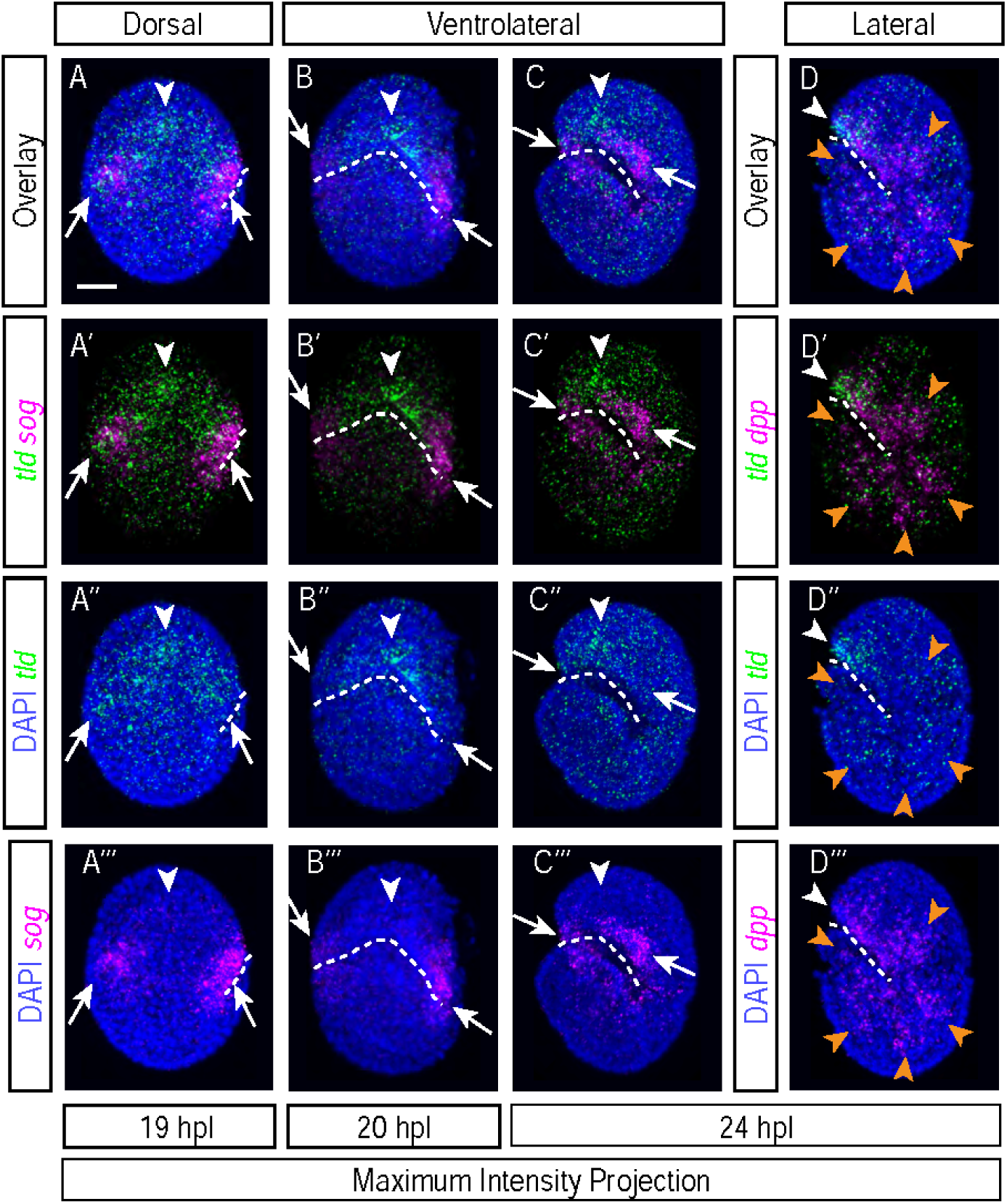
mRNAs of protease tld become more restricted along the anterior-most ridge of the head from 19 to 24 hpl. A-C. Enrichment patterns of *tld* and *sog* mRNAs at 19 hpl (A, n=1 experiment, 6 embryos), 20 hpl (B, n=1 experiment, 7 embryos), and 24 hpl (C, n=4 experiments, 5, 7, 2, and 2 embryos). D. Enrichment patterns of *tld* and *dpp* mRNAs at 24 hpl (n=5 experiments, 13, 2, 1, 2, and 3 embryos). All images are maximum intensity projections. Arrowheads indicate the band of *tld* along the anterior ridge of the embryo. Arrows indicate bands of cells enriched for *sog* on either side of the developing mouth. Orange arrowheads indicate the bands of *dpp* running over each segment. Dorsal view in A. Left ventrolateral view in B-C. Left lateral view in D. Scale bar = 10 μm.

We speculate that, in conjunction with Sog, Tld might facilitate spreading and then release of Sog-bound Dpp either in the region of the developing mouth or internally to the mesodermal precursors – in the endomesodermal pouches of cells surrounding the foregut, which would be consistent with a possibility that Dpp might act to promote mouth development. In addition, Tld is known to function in processing collagens to mediate extracellular matrix (ECM) formation (Vadon-Le Goff et al., 2015). ECM has been shown to be important for pharyngeal morphogenesis in other ecdysozoan animals, such as the nematode *Caenorhabditis elegans* (Rasmussen et al., 2012). Therefore, it is possible that in *H. exemplaris* embryos, Tld plays a role in pharyngeal development independent of BMP.

Based on all of the results examining expression of BMP pathway genes, we conclude that the ligand-encoding gene *dpp* is expressed primarily in lateral ectoderm and in bands near segment boundaries, as well as in the head, and the BMP antagonist-encoding gene *sog* is also expressed in lateral ectoderm, less dorsally than *dpp*, including in the head, where two bands of *sog*-expressing cells are positioned between bands of *dpp*-expressing cells and flanking an anterior band of cells expressing the protease *tld*.

The expression patterns of *doc1*, *eya*, and *dpp* after segmentation of *H. exemplaris* embryos.

These expression patterns left us wondering where BMP and FGF signaling occurs during *H. exemplaris* segmentation. We turned to conserved downstream transcriptional targets of these pathways, *doc1* and *eya,* as likely markers of active signaling. *doc1* and *eya* are conserved targets of BMP signaling in other systems, such as *D. melanogaster* (Dominguez et al., 2016). Additionally, *eya* is known to be a target of FGF signaling (Ahrens and Schlosser, 2005). At ectodermal segmentation (24 hpl), *doc1* was expressed in ectoderm along the dorsal midline of the embryo, running in a band from the posterior tip of the embryo to the top of the head and then spreading into a larger region that did not extend to the anterior tip of the head (Fig. 8 A and B). *doc1* was mostly absent from internal germ layers. The expression of *doc1* in lateral ectoderm is consistent with the hypothesis that *doc1* is a target of Dpp, which has spread from lateral positions up to the dorsal ridge of the embryo, as well as from *dpp*-expressing cells on either side of the development mouth in the head. At ectodermal segmentation (24 hpl), *eya* was expressed in endomesodermal cells and the primordial germ cells and was almost entirely absent from ectoderm (Fig. 8 C and D). Expression of both *doc1* and *eya* appeared non-overlapping with the lateral ectoderm expression of *dpp* (Fig. 8). The expression of *eya* primarily in endomesoderm matches the pattern of *fgf8*-non-expressing, *fgfrl1*-expressing cells that are likely receptive to FGF signaling. We speculate it may mark endomesodermal cells in which FGF signaling is active. The expression of *eya* in primordial germ cells is consistent with the possibility that this gene might be a target of Gbb signaling and that Gbb regulates primordial germ cell development.

**Figure 8.**
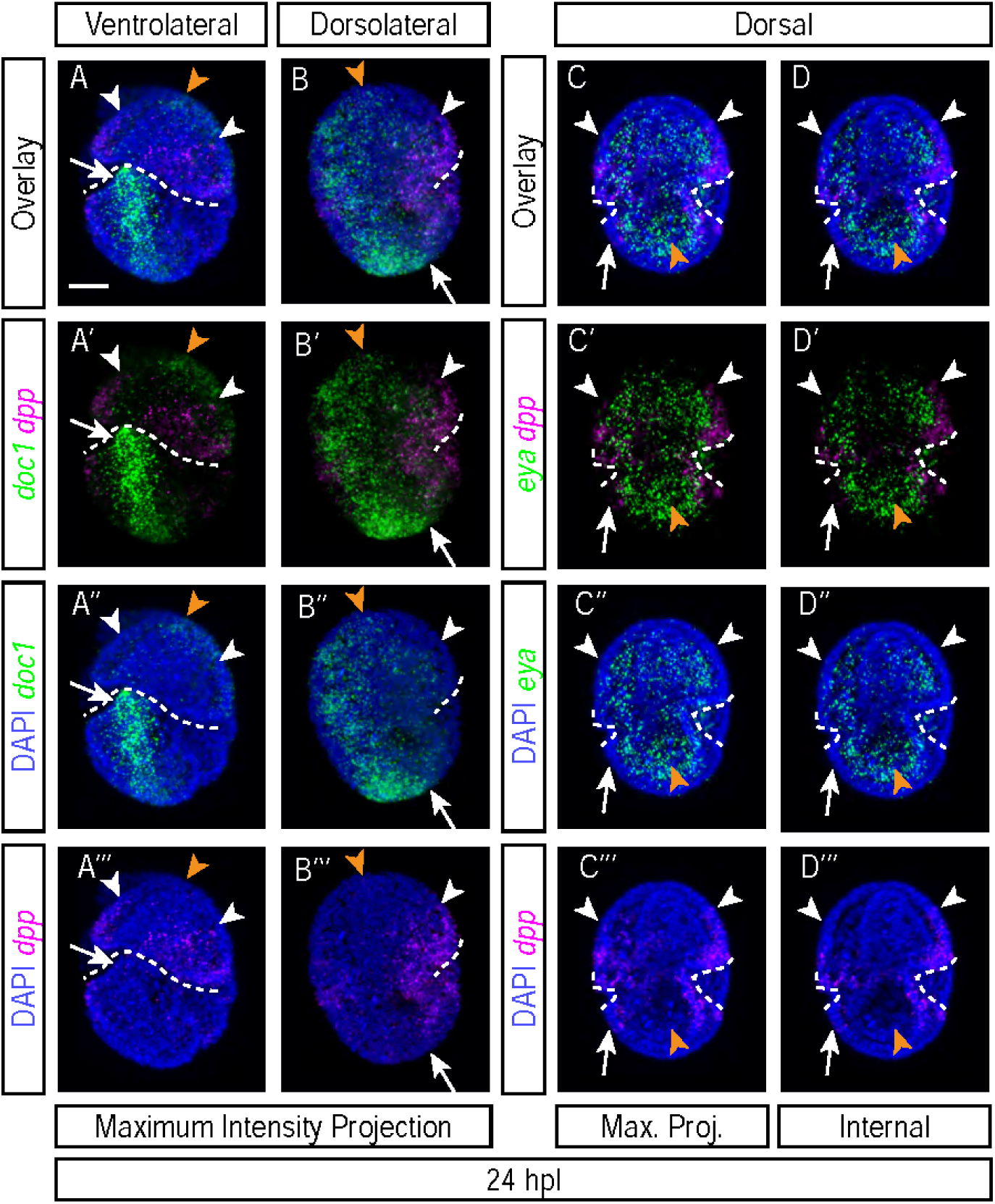
Expression patterns of *doc1, eya,* and *dpp* at ectodermal segmentation stage (24 hpl). A-B. Enrichment patterns of *doc1* and *dpp* mRNAs at 24 hpl (n=4 experiments, 10, 1, 5, and 2 embryos). C-D. Enrichment patterns of *eya* and *dpp* mRNAs at 24 hpl (n=2 experiments, 9 and 2 embryos). A-C. Maximum intensity projections of embryos. D. Projection of internal layers of the embryo in C. Arrowheads in A-D indicate the extend of spread of *dpp* mRNA dorsally. Orange arrowheads in A-B indicate the anterior end of the band of *doc1* along the dorsal midline. Arrows in A-B indicate the posterior end of the band of *doc1* along the dorsal midline. Arrows in C-D indicate the ectodermal layer, which is not enriched for *eya*. Orange arrowheads in C-D indicate the region of the PGCs, which is enriched for *eya*. Left ventrolateral view in A. Right dorsolateral view in B. Dorsal view in C-D. Scale bar = 10 μm.

Finally, between elongation and segmentation stages of *H. exemplaris* embryos, mesoderm forms. Given the known overlap of FGF and BMP in regulating mesoderm in other animals and given that these two pathways exhibit dynamic enrichment patterns between these stages along developing mesoderm, we speculate that FGF and BMP might work together to regulate mesodermal formation and/or fate specification (Row et al., 2018).

## Conclusions

We have detected expression patterns for genes of components of the FGF and BMP signaling pathways during *H. exemplaris* development at segmentation stages (summarized in Fig. 9). We found that expression of the FGF ligand *fgf8* and the FGF receptor *fgfrl1* are enriched in different germ layers between elongation and ectodermal segmentation stages. *fgf8* is expressed in ectoderm at segment posteriors, and *fgfrl1* is broadly expressed in ectoderm and in underlying endomesoderm, but more enriched in the latter. The segmental expression pattern of *fgf8* precedes visible signs of ectodermal segmentation and arises before or concurrently with endomesodermal pouch formation. We also found that the BMP ligand *dpp* and antagonist *sog* are both expressed laterally and that *dpp* is expressed more dorsally than *sog*. Additionally, *sog* is expressed in bands of ectoderm on either side of the developing mouth, and *dpp* is expressed in patches slightly overlapping with and adjacent to these bands of *sog* expression. Finally, *sog* is expressed in a single ventral cell of each endomesodermal pouch, possibly in developing ventral trunk ganglia. *dpp* is expressed in bands running along segment boundaries concomitant with or near the time of ectodermal segmentation. These patterns are apparent by elongation stages and become more elaborate during the morphological process of segmentation.

**Figure 9.**
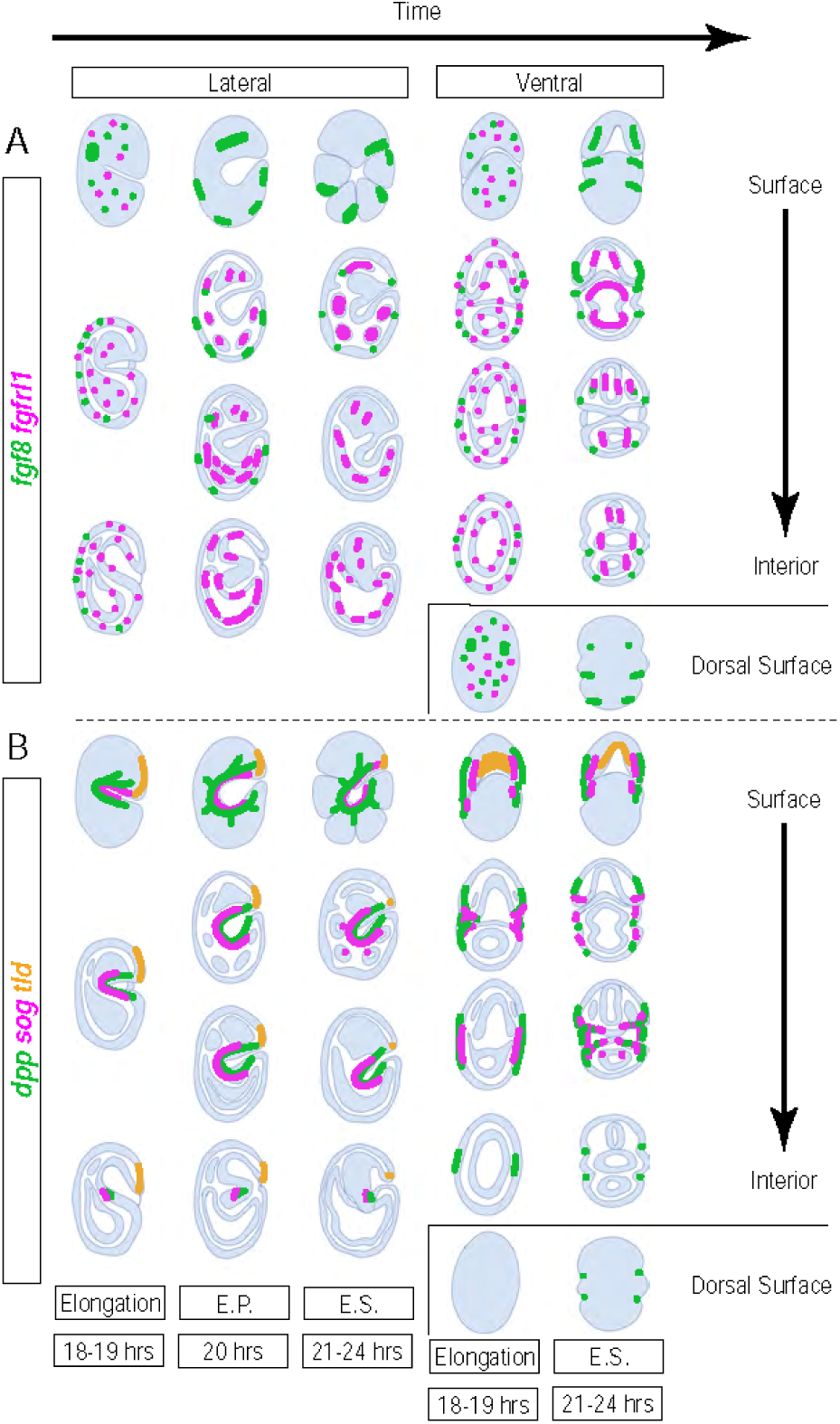
Summary of BMP and FGF pathway gene patterns of enrichment in *H. exemplaris* embryos between elongation and segmentation stages. Drawings representative of enrichment patterns of (A) FGF and (B) BMP signaling pathway genes in *H. exemplaris* embryos at elongation, endomesodermal pouch formation (E.P.), and ectodermal segmentation (E.S.) stages, shown from lateral and ventral views. From top to bottom, images show surface to internal views (and dorsal surface below ventral views). Anterior (A) is up and Posterior (P) is down in all drawings. Dorsal (D) and (V) are to the left and right, respectively, in the lateral views. Representative summaries of expression patterns are shown as follows: in the top embryos (A), *fgf8* in green, *fgfrl1* in magenta; in the bottom embryos (B), *dpp* in green, *sog* in magenta, *tld* in orange.

Based on the expression patterns revealed, we presented hypotheses for the potential roles of FGF and BMP in *H. exemplaris* development (see Table 3 for a more comprehensive summary of hypotheses). Testing these will rely on development of techniques not yet established for this system. Our attempts to use RNAi and chemical inhibitors to FGF (SU5402 and U0126) and BMP signaling (Dorsomorphin, DMH1, and LDN-212854) were unsuccessful (data not shown). Attempting to mark active FGF and BMP signaling, we also stained embryos using cross-reactive anti-pSMAD and anti-dpERK antibodies but did not detect specific signal (data not shown). There has been recent progress towards development of genetic tools for *H. exemplaris* by CRISPR/Cas9-based genome editing of adults with 10-20% efficient genomic deletion and via a vector-based expression system that worked mosaically in adult cells at 70-89% adult transfection efficiency and rare germline transmission (Kumagai et al., 2022; Tanaka et al., 2022). Recently, there has been progress in another tardigrade species, *Ramazzottius varieornatus*, in which CRISPR reagents injected directly into the maternal body cavity exhibited germline transmission with low efficiency (3-4%) (Kondo et al., 2024). We anticipate improvements to these techniques for efficient germline transmission are soon to follow, which would facilitate asking questions about development. Specifically, tools such as transcriptional reporters, as well as inducible protein regulation, will enable temporally specific tests of signaling pathway functions during key stages in development.

Understanding how cell signaling regulates the development of the unique tardigrade body plan will provide new information about how these conserved pathways regulate development of diverse tissue and organismal forms and ultimately how diverse body forms evolved through varied deployment of conserved signaling pathways. Expression analysis among additional tardigrade species and other animal groups will likely be valuable to grasp how nature deploys the same pathways to build widely different or convergently similar animal bodies.

## Experimental Procedures

### Maintaining cultures of *Hypsibius exemplaris*

Cultures were maintained as described previously (Heikes et al., 2023; McNuff, 2018).

### DIC imaging of development

DIC microscopy of embryonic development was performed as previously described (Heikes et al., 2023; Heikes and Goldstein, 2018).

### Gene identification and phylogenetic analyses

Genes were identified as previously described (Heikes et al., 2023). Briefly, genes were identified in a published *Hypsibius exemplaris* transcriptome (Levin et al., 2016) by tBLASTn (Altschul et al., 1990; Gerts et al., 2006; Sayers et al., 2022) using protein sequences of known homologs in *Drosophila melanogaster* from the UCSC genome browser (Kent et al., 2002). A homolog for *fgf8* could not be found in this transcriptome but was identified in another (Yoshida et al., 2017). Top hits were confirmed by reciprocal BLASTp (Sayers et al., 2022) against the *D. melanogaster* transcriptome and further confirmed through comparison of domains to the query sequence using SMART domain analysis tool in normal mode (Letunic et al., 2021; Letunic and Bork, 2018). Phylogenetic reconstructions were produced using alignments of conserved regions (either by manually-curated conserved domains or through the Gblocks program) (Castresana, 2000). Alignments were produced using Neighbor-Joining based MUSCLE alignment of curated protein sequences (Dereeper et al., 2008). Alignments were opened in Jalview to produce EPS files for figures. Alignments were also used to produce maximum likelihood phylogenetic reconstructions. Phylogenetic reconstructions were produced comparing the top hits for each query to known homologs in various species in Mega-X (500 bootstraps), as previously described (Heikes et al., 2023; Kumar et al., 2018). Accession numbers of protein sequences used in maximum likelihood reconstructions and species name abbreviations for all phylogenetic reconstructions are in Table 1.

**Table 1.** Accession numbers used for phylogenetic analysis.

| **Accession Numbers for ML Protein Phylogenetic Reconstructions** |  |  |  |  |  |
| --- | --- | --- | --- | --- | --- |
| **Species Abbrev.** | **FGF8-like** | **Source** | **Sequence ID** | **Translation?** | **Species Name** |
| He | Fgf8 | GenBank<br>PRJNA360553 | BV898_11515_t01 | expasy | *Hypsibius exemplaris* |
| Dm | Ths | UCSC | NP_610701 |  | *Drosophila melanogaster* |
| Ce | Egl-17 | UCSC | NP_508107 |  | *Caenorhabditis elegans* |
| Hv | FGF8 | GenBank | XP_012554564.1 |  | *Hydra vulgaris* |
| Sk | Fgf8 | GenBank | ADB22412.1 |  | *Saccoglossus kowalevskii* |
| Dr | Fgf8 | GenBank | AAB82614.1 |  | *Danio rerio* |
| Hs | Fgf8 | GenBank | NP_149353.1 |  | *Homo sapiens* |
| Bf | Fgf8 | GenBank | XP_035674358.1 |  | *Branchiostoma floridae* |
| Ci | Fgf8/17/18 | GenBank | NP_001027648.1 |  | *Ciona intestinalis* |
| Pf | Fgf8/17/18 | GenBank | AKZ18202.1 |  | *Ptychodera flava* |
| Nv | FGF8a | GenBank | ABN70836.1 |  | *Nematostella vectensis* |
| Tt | Fgf8/17/18 | GenBank | ALS19758.1 |  | *Terebratalia transversa* |
| Na | Fgf8/17/18 | GenBank | ALS19767.1 |  | *Novocrania anomala* |
| Ph | Fgf8/17/18 | GenBank | QUP08116.1 |  | *Phoronopsis harmeri* |
| **Species Abbrev.** | **9/16/20 Family of FGFs** | **Source** | **Sequence ID** | **Translation?** | **Species Name** |
| Dm | Pyr | UCSC | NP_001097275 |  | *Drosophila melanogaster* |
| Dm | Bnl | UCSC | NP_732452 |  | *Drosophila melanogaster* |
| Ce | Let-756 | UCSC | NP_498403 |  | *Caenorhabditis elegans* |
| Nv | FGF9 | GenBank | XP_032228973.1 |  | *Nematostella vectensis* |
| Bf | FGF9 | GenBank | XP_035675604.1 |  | *Branchiostoma floridae* |
| Tt | FGF9/16/20 | GenBank | QUP08106.1 |  | *Terebratalia transversa* |
| Na | FGF9/16/20 | GenBank | QUP08110.1 |  | *Novocrania anomala* |
| Ph | FGF9/16/20 | GenBank | QUP08115.1 |  | *Phoronopsis harmeri* |
| Hv | FGF9 | GenBank | AND74489.1 |  | *Hydra vulgaris* |
| Ci | FGF9/16/20 | GenBank | NP_001027649.1 |  | *Ciona intestinalis* |
| Hs | FGF9 | GenBank | NP_002001.1 |  | *Homo sapiens* |
| Dr | FGF9 | GenBank | XP_009302933.1 |  | *Danio rerio* |
| Sk | FGF9 | GenBank | NP_001161538 |  | *Saccoglossus kowalevskii* |
| **Species Abbrev.** | **FGF Receptor-like** | **Source** | **Sequence ID** | **Translation?** | **Species Name** |
| He | FGFRL1 | Levin et al., 2016 transcriptome | He_tr_04670 | expasy | *Hypsibius exemplaris* |
| He | FGFRL2 | Levin et al., 2016 transcriptome | He_tr_03639 | expasy | *Hypsibius exemplaris* |
| Dm | Htl | UCSC | NP_732287 |  | *Drosophila melanogaster* |
| Dm | Btl | UCSC | NP_729956 |  | *Drosophila melanogaster* |
| Ce | Egl-15 | UCSC | NP_001369987 |  | *Caenorhabditis elegans* |
| Nv | FGFRa | GenBank | XP_048584433.1 |  | *Nematostella vectensis* |
| Nv | FGFRb | GenBank | XP_048585038.1 |  | *Nematostella vectensis* |
| Na | FGFR1 | GenBank | QUP08111.1 |  | *Novocrania anomala* |
| Tt | FGFR | GenBank | QUP08107.1 |  | *Terebratalia transversa* |
| Ph | FGFR | GenBank | QUP08114.1 |  | *Phoronopsis harmeri* |
| Hs | FGFR1 | GenBank | AAH15035.1 |  | *Homo sapiens* |
| Hs | FGFR2 | GenBank | CAA96492.1 |  | *Homo sapiens* |
| Hs | FGFR3 | GenBank | XP_047305776.1 |  | *Homo sapiens* |
| Hs | FGFR4 | GenBank | NP_001341913.1 |  | *Homo sapiens* |
| Bf | FGFR | GenBank | XP_035673320.1 |  | *Branchiostoma floridae* |
| **Species Abbrev.** | **Non-FGFR RTKs** | **Source** | **Sequence ID** | **Translation?** | **Species Name** |
| He | VEGFR | Yanai | He_tr_02285 | expasy | *Hypsibius exemplaris* |
| Dm | PVR | UCSC | NP_723365 |  | *Drosophila melanogaster* |
| Ce | VER-1 | UCSC | NP_497162 |  | *Caenorhabditis elegans* |
| Ce | VER-3 | UCSC | NP_509836 |  | *Caenorhabditis elegans* |
| Ce | VER-4 | UCSC | NP_509835 |  | *Caenorhabditis elegans* |
| Hs | VGFR1 | GenBank | NP_002010.2 |  | *Homo sapiens* |
| Hs | VGFR2 | GenBank | NP_002244.1 |  | *Homo sapiens* |
| Hs | VGFR3 | GenBank | NP_891555.2 |  | *Homo sapiens* |
| Bf | VEGFR | GenBank | XP_035682136.1 |  | *Branchiostoma floridae* |
| **Species Abbrev.** | **Snail** | **Source** | **Sequence ID** | **Translation?** | **Species Name** |
| He | Snail | Levin et al., 2016 transcriptome | Hduj_tr_05604 | expasy | *Hypsibius exemplaris* |
| Dm | Snail | UCSC | NP_476732 |  | *Drosophila melanogaster* |
| Dm | Escargot | UCSC | NP_476600 |  | *Drosophila melanogaster* |
| Nv | SnaA | GenBank | AAR24456.1 |  | *Nematostella vectensis* |
| Nv | SnaB | GenBank | AAR24457.1 |  | *Nematostella vectensis* |
| Tc | Snail | GenBank | XP_971329.1 |  | *Tribolium castaneum* |
| Hv | Snail | GenBank | XP_002162096.1 |  | *Hydra vulgaris* |
| Ci | Snail | GenBank | AAB61226.1 |  | *Ciona intestinalis* |
| Pf | Snail | GenBank | AXA20416.1 |  | *Ptychodera flava* |
| Hs | SNAI1 | GenBank | NP_005976.2 |  | *Homo sapiens* |
| Bf | Snail | GenBank | AAC35351.1 |  | *Branchiostoma floridae* |
| **Species Abbrev.** | **Scratch** | **Source** | **Sequence ID** | **Translation?** | **Species Name** |
| He | Scratch | Levin et al., 2016 transcriptome | Hduj_tr_06310 | expasy | *Hypsibius exemplaris* |
| Dm | Scrt | UCSC | NP_523911.2 |  | *Drosophila melanogaster* |
| Ce | CES-1 | UCSC | NP_492338 |  | *Caenorhabditis elegans* |
| Nv | Scratch2 | GenBank | XP_032237161.1 |  | *Nematostella vectensis* |
| Tc | Scratch1 | GenBank | EFA13195.2 |  | *Tribolium castaneum* |
| Tc | Scratch2 | GenBank | EFA13196.1 |  | *Tribolium castaneum* |
| Hs | Scratch1 | GenBank | NP_112599.2 |  | *Homo sapiens* |
| Hs | Scratch2 | GenBank | NP_149120.1 |  | *Homo sapiens* |
| Bf | Scratch1 | GenBank | XP_035679385.1 |  | *Branchiostoma floridae* |
| Bf | Scratch2 | GenBank | XP_035679386.1 |  | *Branchiostoma floridae* |
| **Species Abbrev.** | **Dpp/BMP2/4** | **Source** | **Sequence ID** | **Translation?** | **Species Name** |
| He | Dpp | Levin et al., 2016 transcriptome | He_tr_08636 | expasy | *Hypsibius exemplaris* |
| Dm | Dpp | UCSC | NP_477311 |  | *Drosophila melanogaster* |
| Dm | Scw | UCSC | NP_524863 |  | *Drosophila melanogaster* |
| Mm | BMP2 | GenBank | NP_031579.2 |  | *Mus musculus* |
| Mm | BMP4 | GenBank | NP_001303289.1 |  | *Mus musculus* |
| Nv | BMP2/4 | GenBank | AAR13362.1 |  | *Nematostella vectensis* |
| Tc | Dpp | GenBank | NP_001034540.1 |  | *Tribolium* |
|  |  |  |  |  | *castaneum* |
| **Species Abbrev.** | **Gbb/BMP5/7/8** | **Source** | **Sequence ID** | **Translation?** | **Species Name** |
| He | Gbb | Levin et al., 2016 transcriptome | He_tr_00999 | expasy | *Hypsibius exemplaris* |
| Dm | Gbb | UCSC | NP_477340 |  | *Drosophila melanogaster* |
| Mm | BMP5 | GenBank | NP_031581.2 |  | *Mus musculus* |
| Mm | BMP7 | GenBank | NP_031583.2 |  | *Mus musculus* |
| Mm | BMP8B | GenBank | NP_031585.2 |  | *Mus musculus* |
| Mm | BMP8A | GenBank | NP_031584.1 |  | *Mus musculus* |
| Nv | BMP5/7/8 | GenBank | XP_032219054.1 |  | *Nematostella vectensis* |
| Tc | Gbb | GenBank | NP_001107813.1 |  | *Tribolium castaneum* |
| **Species Abbrev.** | **Other BMPs** | **Source** | **Sequence ID** | **Translation?** | **Species Name** |
| Mm | BMP3 | GenBank | NP_775580.1 |  | *Mus musculus* |
| Mm | BMP6 | GenBank | NP_031582.1 |  | *Mus musculus* |
| Mm | BMP9/GDF2 | GenBank | NP_062379.3 |  | *Mus musculus* |
| Mm | BMP10 | GenBank | NP_033886.2 |  | *Mus musculus* |
| **Species Abbrev.** | **Sog/Chordin** | **Source** | **Sequence ID** | **Translation?** | **Species Name** |
| He | sog | Levin et al., 2016 transcriptome | He_tr_06660 | expasy | *Hypsibius exemplaris* |
| Dm | sog | UCSC | NP_001259578 |  | *Drosophila melanogaster* |
| Mm | chordin | NCBI_GenBank | AAD19895.1 |  | *Mus musculus* |
| Xl | chordin | NCBI_GenBank | AAC42222.1 |  | *Xenopus laevis* |
| Tc | sog | NCBI_GenBank | ABF22614.1 |  | *Tribolium castaneum* |
| Am | sog | NCBI_GenBank | XP_006570089.1 |  | *Apis mellifera* |
| Hs | chordin | NCBI_GenBank | AAG35767.1 |  | *Homo sapiens* |
| Ci | chordin | NCBI_GenBank | XP_026690585.1 |  | *Ciona intestinalis* |
| Nv | chordin | NCBI_GenBank | ABC88373.1 |  | *Nematostella vectensis* |
| **Species Abbrev.** | **Tkv/Type I receptor** | **Source** | **Sequence ID** | **Translation?** | **Species Name** |
| He | Tkv | Levin et al., 2016 transcriptome | He_tr_10719 | expasy | *Hypsibius exemplaris* |
| Dm | Tkv | UCSC | NP_787989 |  | *Drosophila melanogaster* |
| Gb | Tkv | NCBI_GenBank | AHJ59839.1 |  | *Gryllus bimaculatus* |
| Tc | Tkv | NCBI_GenBank | EFA09250.1 |  | *Tribolium castaneum* |
| Mm | BMPR1a | NCBI_GenBank | XP_036014306.1 |  | *Mus musculus* |
| Nv | BMPR1b | NCBI_GenBank | XP_001633896.2 |  | *Nematostella vectensis* |
| **Species Abbrev.** | **Punt/Type II receptor** | **Source** | **Sequence ID** | **Translation?** | **Species Name** |
| He | Punt | Levin et al., 2016 transcriptome | He_tr_12285 | expasy | *Hypsibius exemplaris* |
| Dm | Punt | UCSC | NP_001262574 |  | *Drosophila melanogaster* |
| Tc | Punt | NCBI_GenBank | EEZ97734.1 |  | *Tribolium castaneum* |
| Mm | BMPR2 | NCBI_GenBank | EDL00137.1 |  | *Mus musculus* |
| Nv | BMPR2 | NCBI_GenBank | XP_048588435.1 |  | *Nematostella vectensis* |
| **Species Abbrev.** | **Sax/Type II receptor** | **Source** | **Sequence ID** | **Translation?** | **Species Name** |
| He | Sax | Levin et al., 2016 transcriptome | He_tr_01951 | expasy | *Hypsibius exemplaris* |
| He | Babo | Levin et al., 2016 transcriptome | He_tr_12110 | expasy | *Hypsibius exemplaris* |
| Dm | Sax | UCSC | NP_523652 |  | *Drosophila melanogaster* |
| Dm | Babo | UCSC | NP_477000 |  | *Drosophila melanogaster* |
| Tc | Babo | NCBI_GenBank | KYB28937.1 |  | *Tribolium castaneum* |
| Tc | Sax | NCBI_GenBank | CDW20669.1 |  | *Tribolium castaneum* |
| Dm | Wit | NP_524692 | UCSC |  | *Drosophila melanogaster* |
| Tc | Wit | EFA07475.1 | NCBI_GenBank |  | *Tribolium castaneum* |
| **Species Abbrev.** | **Tld** | **Source** | **Sequence ID** | **Translation?** | **Species Name** |
| He | Tld | Levin et al., 2016 transcriptome | Hduj_tr_05721 | expasy | *Hypsibius exemplaris* |
| Dm | Tld | UCSC | NP_524487 |  | *Drosophila melanogaster* |
| Dr | Tld | NCBI_GenBank | AAC60304.1 |  | *Danio rerio* |
| Gg | Tld | NCBI_GenBank | NP_990034.2 |  | *Gallus gallus* |
| Tc | Tld | NCBI_GenBank | KYB28517.1 |  | *Tribolium castaneum* |
| Am | Tld | NCBI_GenBank | XP_006567026.2 |  | *Apis mellifera* |
| Hs | Tld | NCBI_GenBank | KAI4027581.1 |  | *Homo sapiens* |
| Ci | Tld | NCBI_GenBank | NP_001071840.1 |  | *Ciona intestinalis* |
| Nv | Tld | NCBI_GenBank | XP_001633846.2 |  | *Nematostella vectensis* |
| **Species Abbrev.** | **Doc1** | **Source** | **Sequence ID** | **Translation?** | **Species Name** |
| He | Doc1 | Levin et al., 2016 transcriptome | HD_v1_tr_01406 | expasy | *Hypsibius exemplaris* |
| Dm | Doc1 | UCSC | NP_648283 |  | *Drosophila melanogaster* |
| Tc | Doc | NCBI_GenBank | EFA10735.1 |  | *Tribolium castaneum* |
| Mm | Tbx6 | NCBI_GenBank | AAT72924.1 |  | *Mus musculus* |
| Xl | Tbx6 | NCBI_GenBank | ABC75836.1 |  | *Xenopus laevis* |
| Xl | VegT | NCBI_GenBank | AAB93301.1 |  | *Xenopus laevis* |
| Mm | Brachyury | NCBI_GenBank | AAI20808.1 |  | *Mus musculus* |
| Mm | Tbx2 | NCBI_GenBank | AAC52697.1 |  | *Mus musculus* |
| Mm | Tbx20 | NCBI_GenBank | NP_919239.1 |  | *Mus musculus* |
| He | Omb | Levin et al.,<br>2016<br>transcriptome | Hduj_tr_06618 | expasy | *Hypsibius*<br>*exemplaris* |
| Dm | Omb | UCSC | NP_525070 |  | *Drosophila*<br>*melanogaster* |
| **Species**<br>**Abbrev.** | **Eya** | **Source** | **Sequence ID** | **Translation?** | **Species Name** |
| He | Eya | Levin et al.,<br>2016<br>transcriptome | HD_v1_tr_10204 | expasy | *Hypsibius*<br>*exemplaris* |
| Dm | Eya | UCSC | NP_723188 |  | *Drosophila*<br>*melanogaster* |
| Ce | Eya-1 | UCSC | NP_001367120 |  | *Caenorhabditis*<br>*elegans* |
| Mm | Eya1 | NCBI_GenBank | NP_001297388.1 |  | *Mus musculus* |
| Dr | Eya1 | NCBI_GenBank | AAI54188.1 |  | *Danio rerio* |
| Hs | Eya1 | NCBI_GenBank | AAI21799.1 |  | *Homo sapiens* |
| Nv | Eya | NCBI_GenBank | XP_032233587.1 |  | *Nematostella*<br>*vectensis* |
| At | Eya | NCBI_GenBank | NP_565803.1 |  | *Arabidopsis*<br>*thaliana* |

### Cloning

Genes were cloned as previously described (Heikes et al., 2023; Smith et al., 2016). Primers were designed using NCBI PrimerBLAST (Ye et al., 2012) to amplify genes by nested PCR. Primers are listed in Table 2.

**Table 2.**
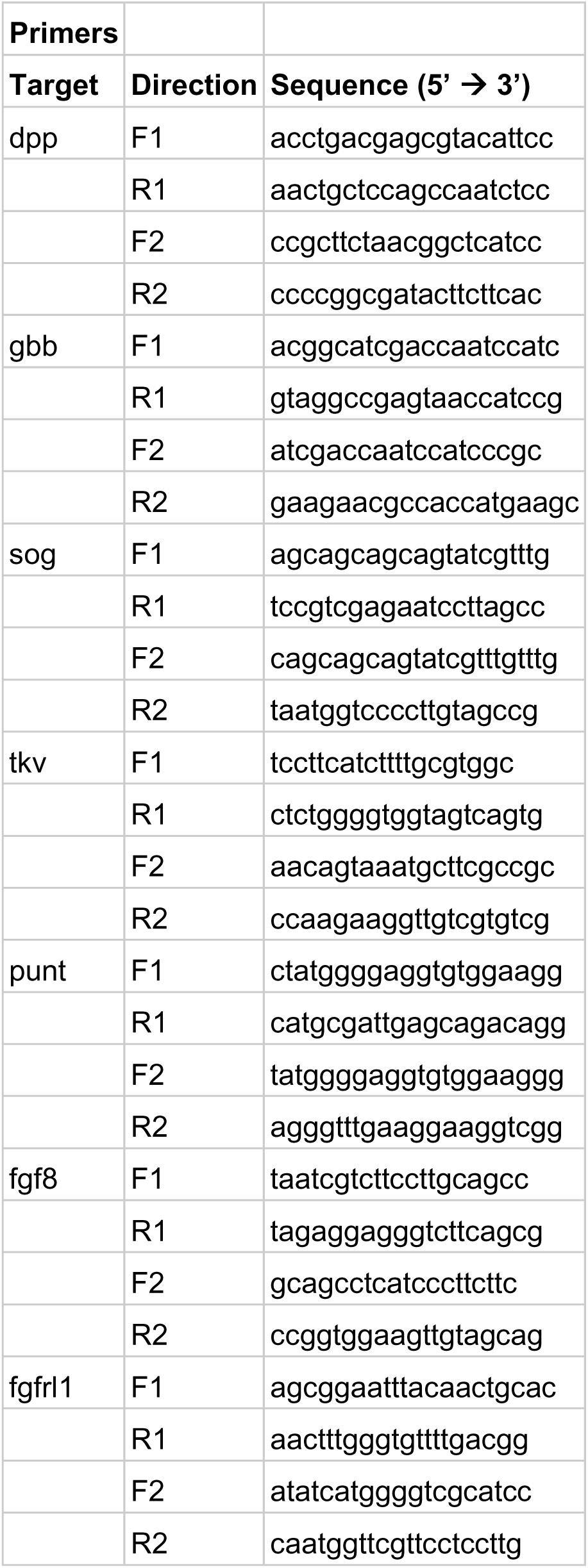
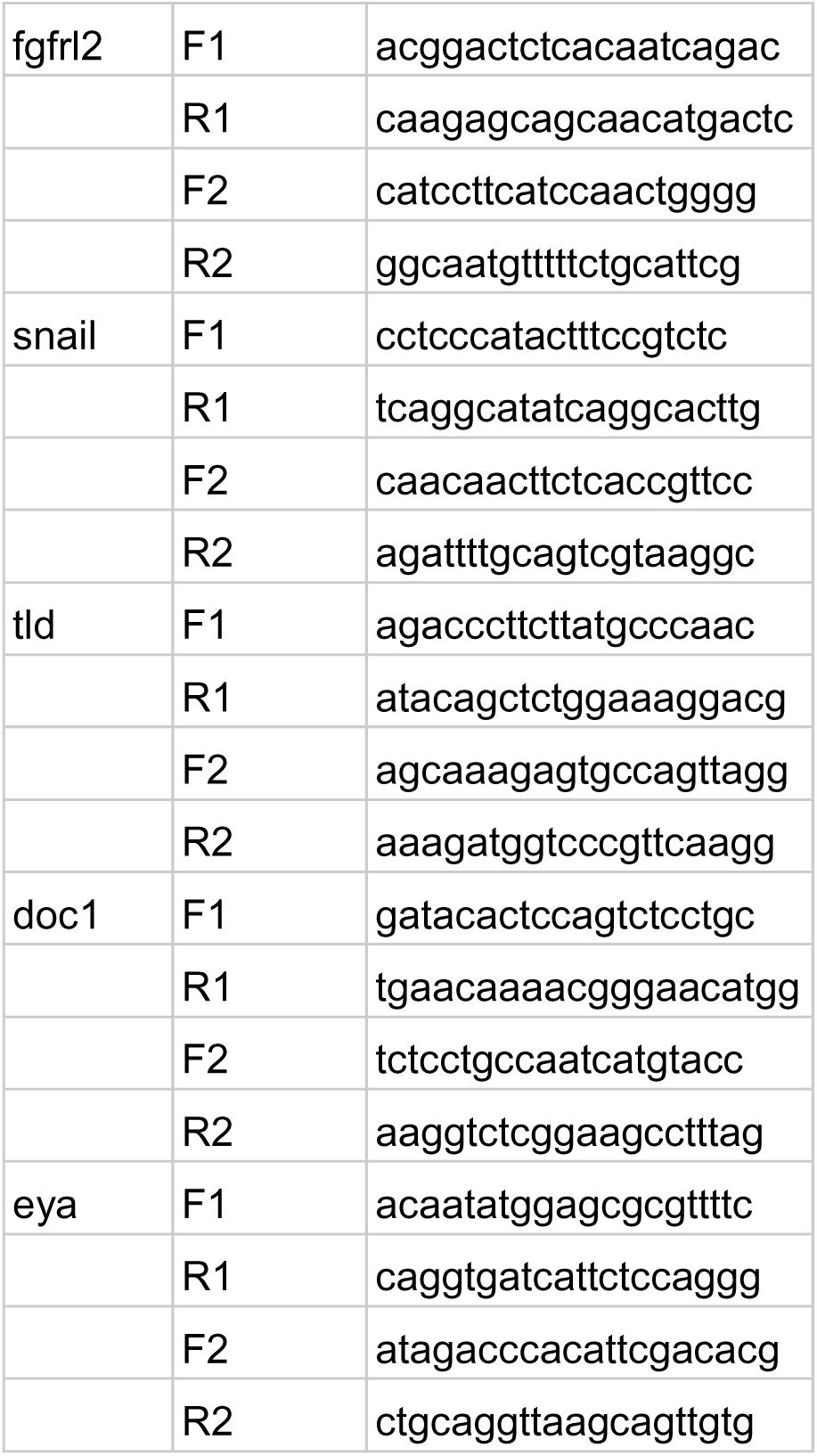
Primers used for cloning.

**Table 3.**
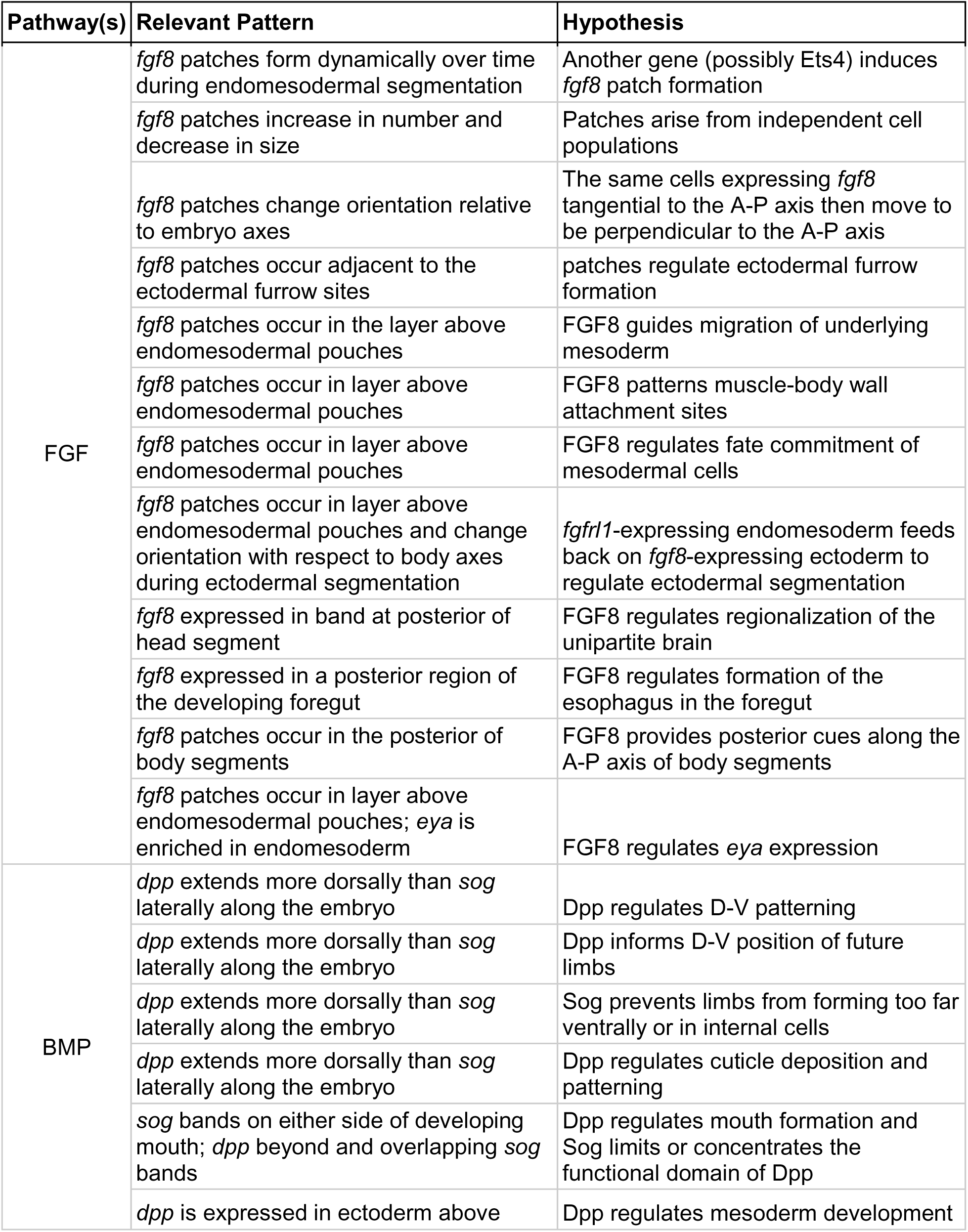
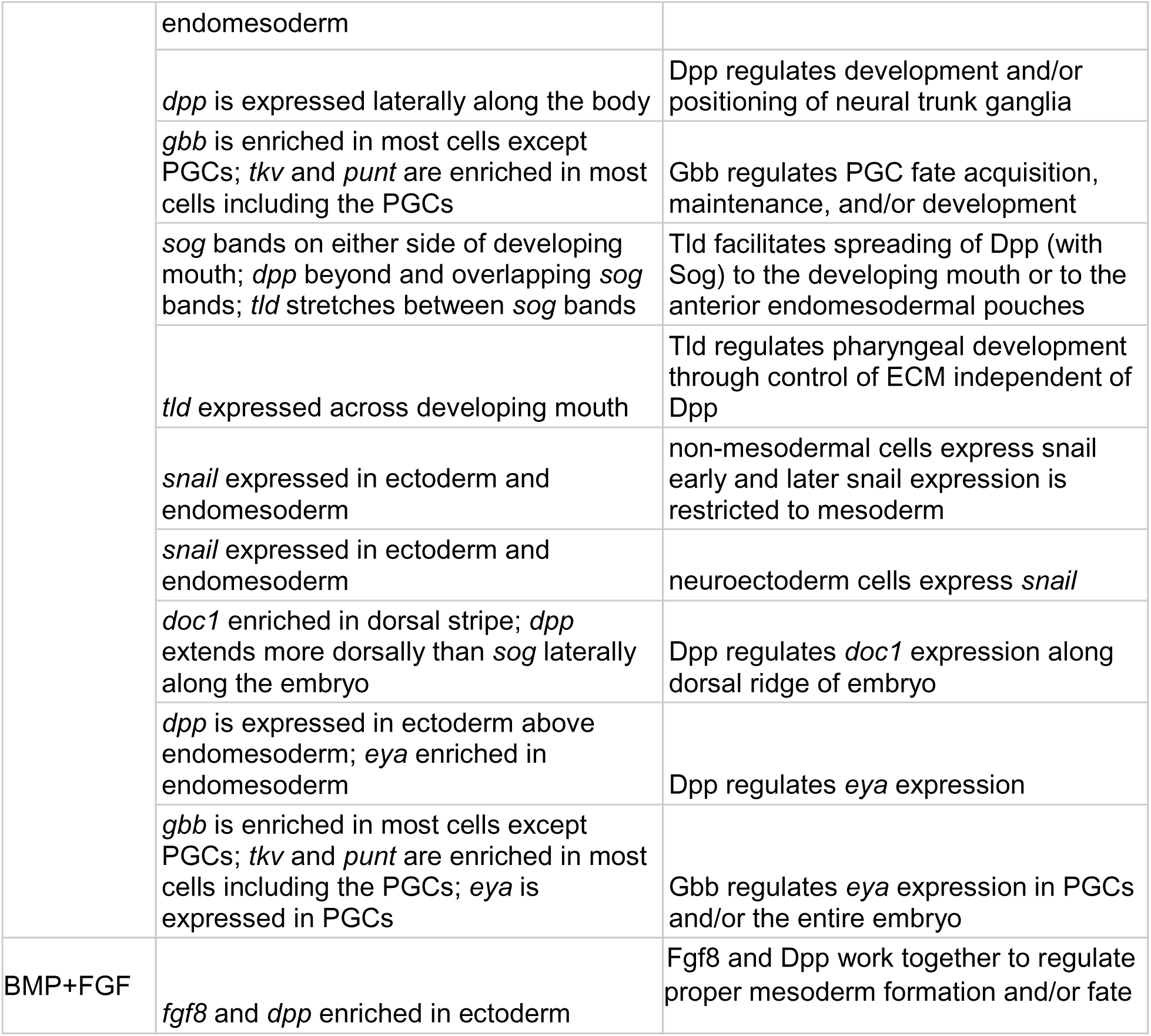
Some hypotheses of FGF and BMP Functions.

### Probe synthesis

RNA probes were synthesized as previously described (Heikes et al., 2023; Smith, 2018; Smith et al., 2016). Probes were labelled with either the DIG RNA labeling mix (Roche, Sigma product # 11277073910) or the Fluorescein RNA labeling mix (Roche, Sigma product # 11685619910) to facilitate double staining.

### Fluorescence *in situ* hybridization

Fluorescence *in situ* hybridization (FISH) was performed using the Tyramide Signal Amplification kit from Akoya Biosciences, as previously described (Heikes et al., 2023), with the addition of a second round of antibody and signal amplification for double FISH. The added steps for double FISH performed were as follows, starting with the end of the tyramide signal amplification (steps 37-42) in the protocol previously published (Heikes et al., 2023). Embryos were washed in pre-heated Solution X for 20 minutes at 60°C (final concentrations: 50% formamide, 2x SSC, 1% SDS, diluted in DEPC-treated water) to inactivate peroxidase activity from the anti-DIG-POD antibody. Embryos were then washed four times quickly in 0.5x PBTw at room temperature. The protocol was then repeated starting from step 34 (Heikes et al., 2023) through to the end of the protocol, with the exception that anti-FITC-POD antibody was used to detect FITC-labelled RNA probes, instead of the anti-DIG-POD antibody used in the first round of detection. For both antibody incubations, a concentration of 1:500 antibody:blocking buffer was used. DIG-labelled probes were amplified with Cy3 amplification reagent (Akoya product # NEL744001KT) and FITC-labelled probes were amplified with Cy5 amplification reagent (Akoya product # NEL745001KT).

### Fluorescence microscopy

FISH-stained embryos were imaged on a Zeiss LSM 710 microscope with a Plan-Neofluor 100x/1.3 oil Iris objective or a Zeiss LSM 880 microscopy with fast Airyscan detector and a Plan-Apochromat 63x/1.4 oil DIC objective. DAPI signal was excited with a 405 nm laser and collected between 411 and 543 nm. Cy3 signal was excited with a 560 nm laser and collected between 561 and 620 nm. Cy5 signal was excited with a 633 nm laser and collected between 639 and 750 nm. Images were opened in FIJI or Zen Black edition for image analysis. Minimum and Maximum displayed values were linearly adjusted in FIJI or Zen Black edition, and images were exported to scale in PNG format for cropping and annotation. Figures were made in Adobe Illustrator. Scale bars were added in Illustrator by pixel scale. DAPI stained DNA signal is displayed as blue in all figures. Cy3-amplified DIG-labelled probe signal is displayed in green in all figures, except for single-channel images, which are displayed in grayscale. Cy5-amplified FITC-labelled probe signal is displayed as magenta in all figures. Figure images are representative of multiple samples across independent rounds of staining, with sample numbers listed in each legend. Anterior is up in all images. Large dashed lines in figures indicate the boundary between the anterior and posterior of embryos.

### Curating diagrams of development

Diagrams of development were hand-drawn on a Samsung Galaxy S8 tablet in the Concepts app, using fluorescence images to trace outlines and expression patterns.

## Acknowledgements

We thank members of the Goldstein lab, past and present, for ideas and promoting a supportive environment conducive to research with an emerging model. In particular, we thank Frank Smith and members of the Smith lab for scientific discussions and for discussing unpublished results freely. We thank the UNC Biology Department, especially Tony Purdue and Nat Prunet for technical assistance in the UNC Biology microscopy core. We thank Dr. Amy Maddox, Dr. Amy Gladfelter, Dr. Greg Wray, and Dr. Christina Burch for helpful feedback on the manuscript and/or scientific discussions.

---

**Supplementary Figure 1.**
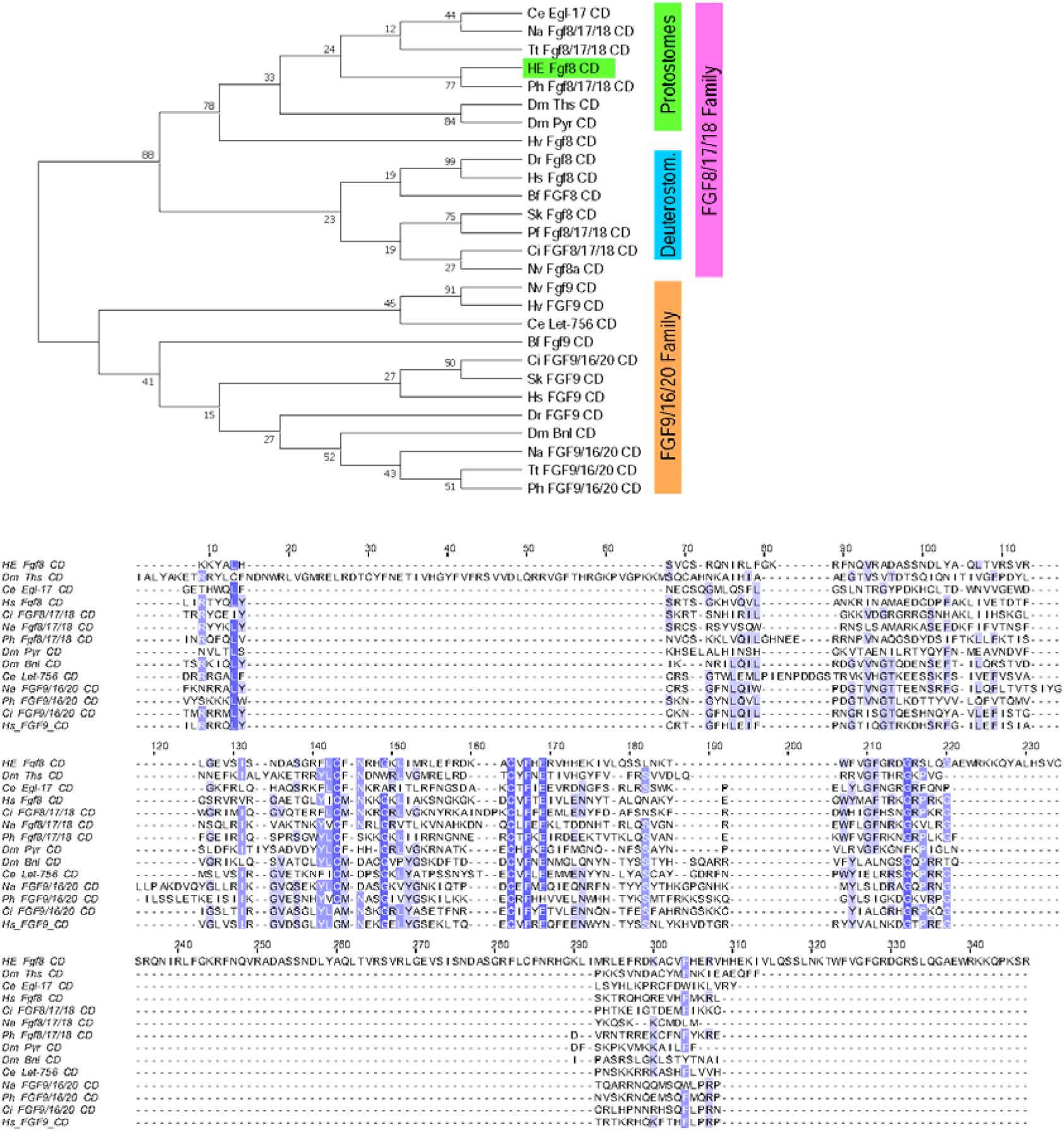
Identification of an FGF8 homolog in *H. exemplaris*. Above: Maximum Likelihood phylogenetic reconstruction of FGF8 and related amino acid sequences. Branch support out of 100 is given at each node. *H. exemplaris* sequences are highlighted. Protein families are indicated by colored bars to the right of the tree. Deuterostom = Deuterostomes. Below: Alignment used for phylogenetic reconstruction from conserved domain among FGF homologs from several species. Genes include 2-letter prefixes indicating species listed in Table 1.

**Supplementary Figure 2.**
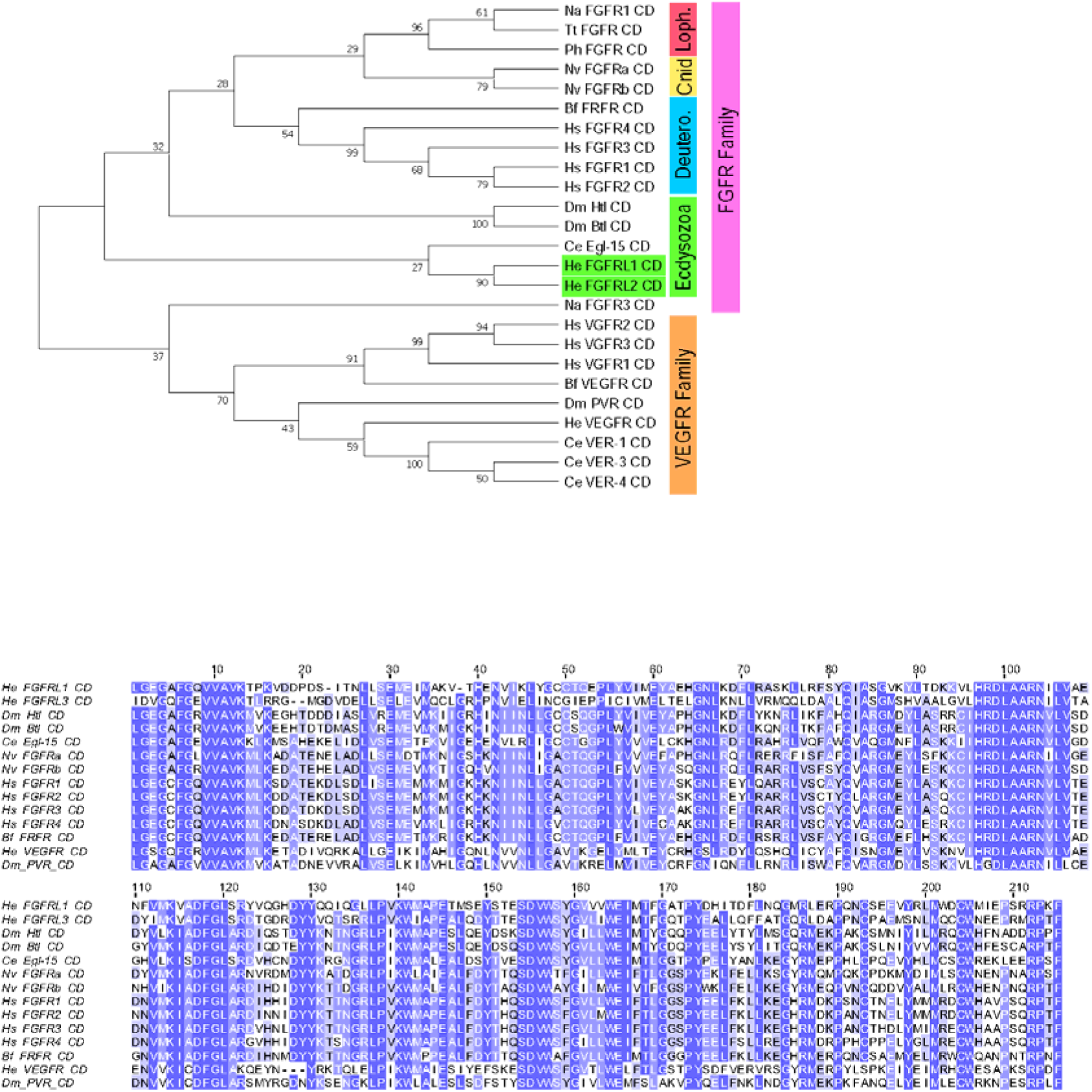
Identification of FGF Receptor homologs in *H. exemplaris*. Above: Maximum Likelihood phylogenetic reconstruction of FGF receptors and related amino acid sequences. Branch support out of 100 is given at each node. *H. exemplaris* sequences are highlighted. Protein families are indicated by colored bars to the right of the tree. Cnid = Cnidaria; Loph. = Lophotrocozoa; Deutero = Deuterostomes. Below: Alignment used for phylogenetic reconstruction from conserved domain among FGF receptor homologs from several species. Genes include 2-letter prefixes indicating species listed in Table 1.

**Supplementary Figure 3.**
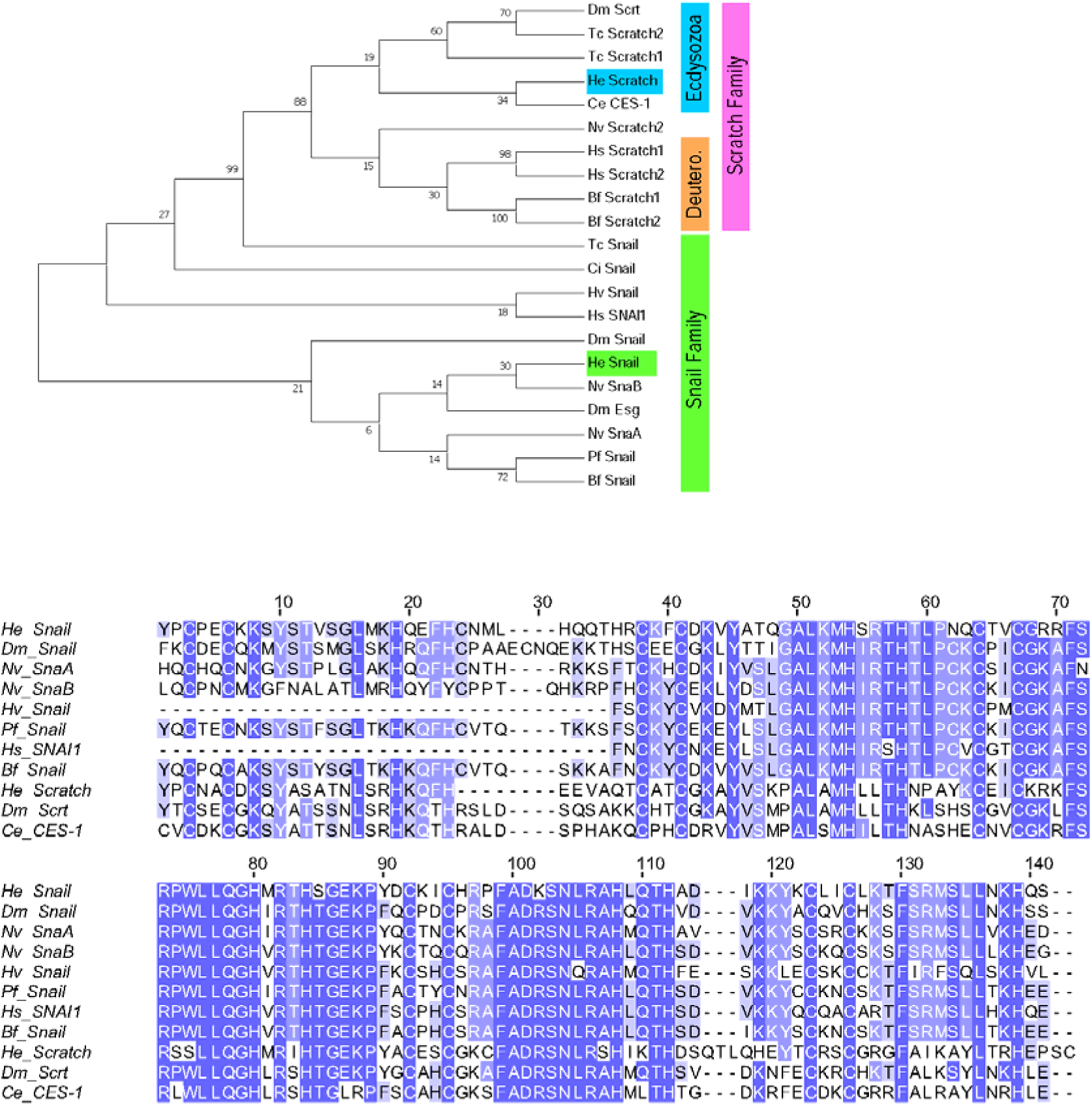
Identification of a Snail homolog in *H. exemplaris*. Above: Maximum Likelihood phylogenetic reconstruction of Snail and related amino acid sequences. Branch support out of 100 is given at each node. *H. exemplaris* sequences are highlighted. Protein families are indicated by colored bars to the right of the tree. Deutero = Deuterostomes. Below: Alignment used for phylogenetic reconstruction from conserved region among Snail homologs from several species. Genes include 2-letter prefixes indicating species listed in Table 1.

**Supplementary Figure 4.**
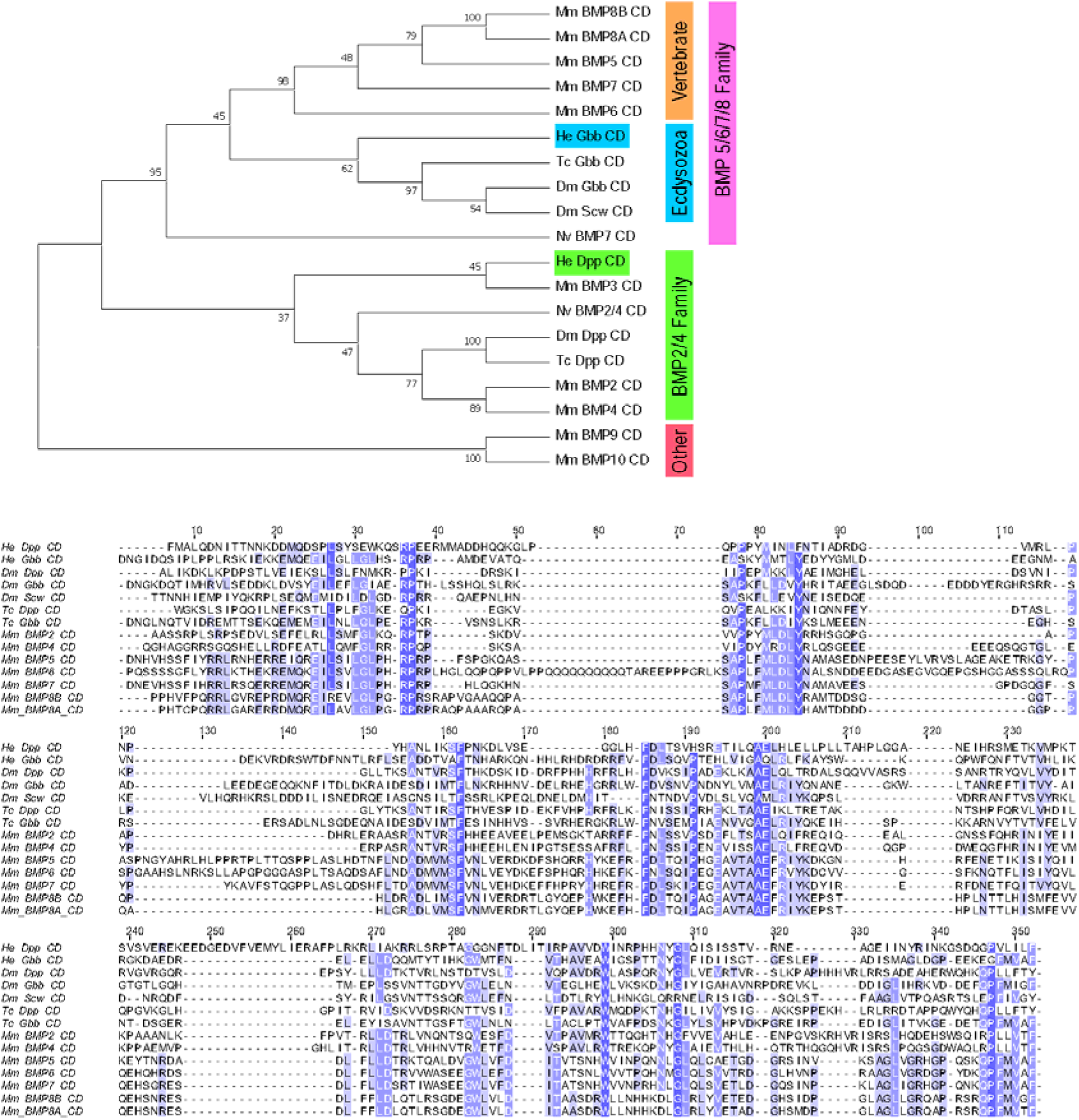
Identification of BMP ligand homologs in *H. exemplaris*. Above: Maximum Likelihood phylogenetic reconstruction of BMP ligands and related amino acid sequences. Branch support out of 100 is given at each node. *H. exemplaris* sequences are highlighted. Protein families are indicated by colored bars to the right of the tree. Below: Alignment used for phylogenetic reconstruction from conserved domain among BMP homologs from several species. Genes include 2-letter prefixes indicating species listed in Table 1.

**Supplementary Figure 5.**
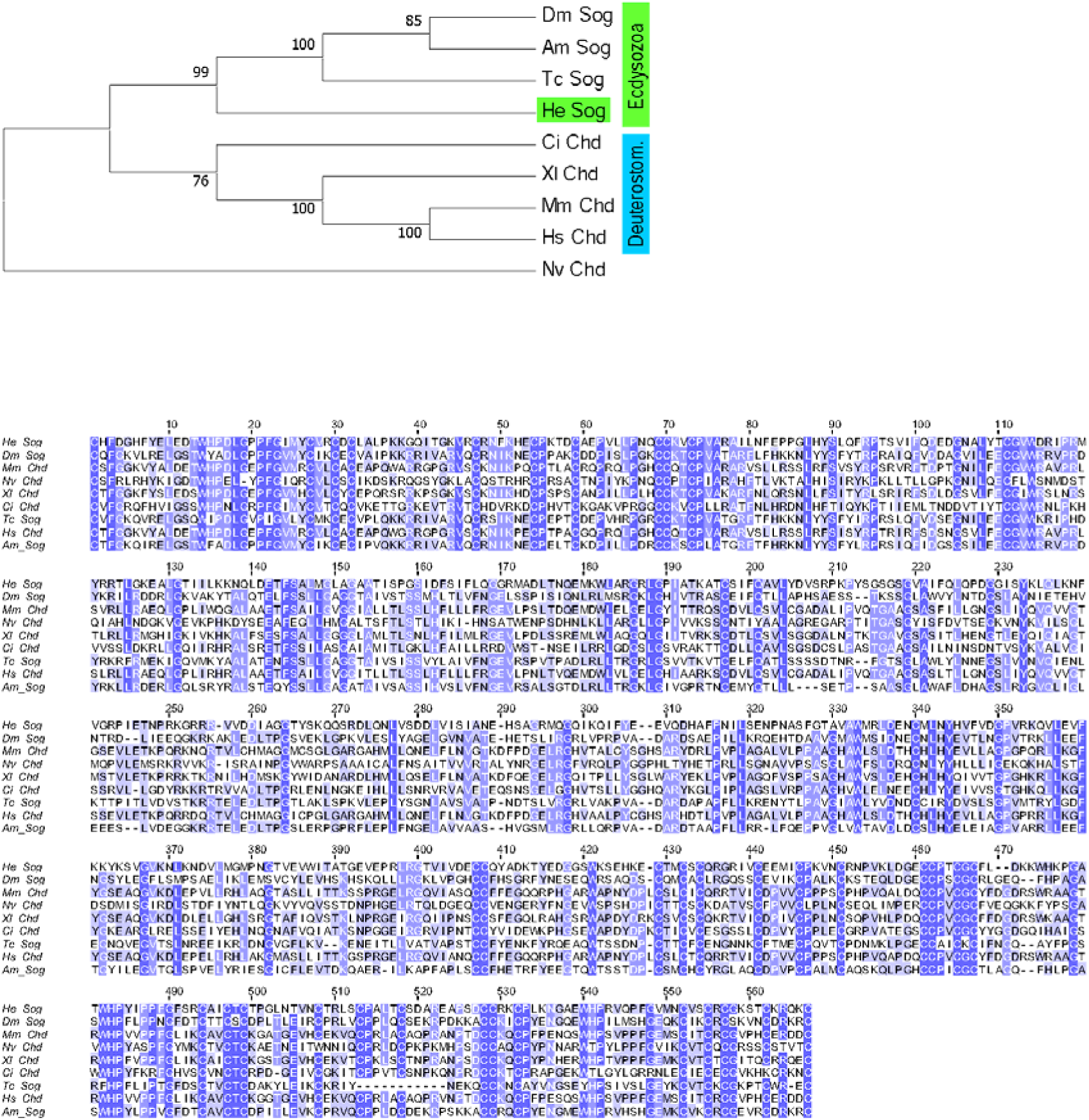
Identification of a Sog homolog in *H. exemplaris*. Above: Maximum Likelihood phylogenetic reconstruction of Sog/Chordin and related amino acid sequences. Branch support out of 100 is given at each node. *H. exemplaris* sequences are highlighted. Protein families are indicated by colored bars to the right of the tree. Deuterostom = Deuterostomes. Below: Alignment used for phylogenetic reconstruction from conserved region among Sog/Chordin homologs from several species. Genes include 2-letter prefixes indicating species listed in Table 1.

**Supplementary Figure 6.**
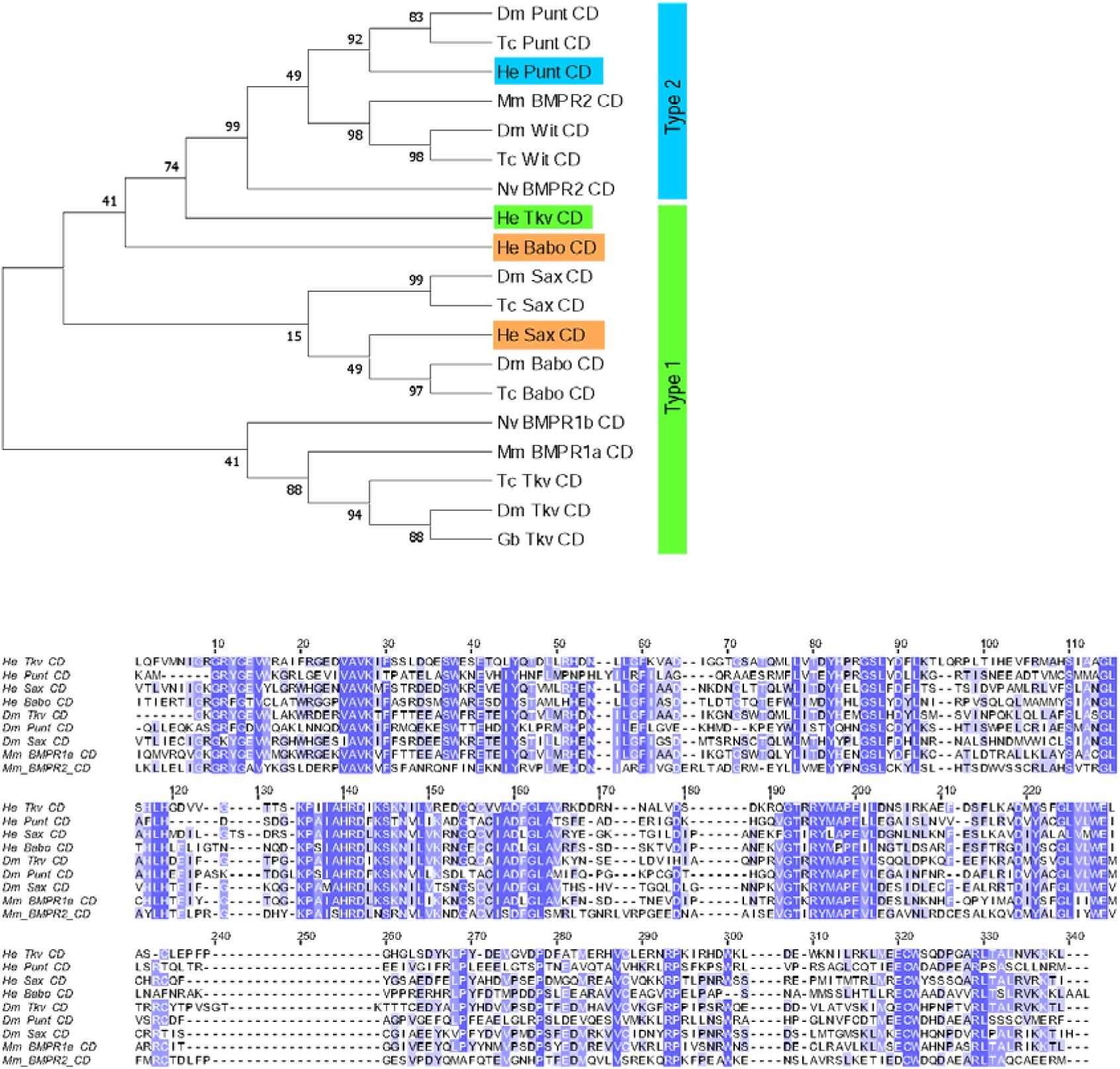
Identification of BMP Type I and II Receptor homologs in *H. exemplaris*. Above: Maximum Likelihood phylogenetic reconstruction of BMP receptors Type 1 and 2 and related amino acid sequences. Branch support out of 100 is given at each node. *H. exemplaris* sequences are highlighted. Protein families are indicated by colored bars to the right of the tree. Below: Alignment used for phylogenetic reconstruction from conserved domain among BMP receptor homologs from several species. Genes include 2-letter prefixes indicating species listed in Table 1.

**Supplementary Figure 7.**
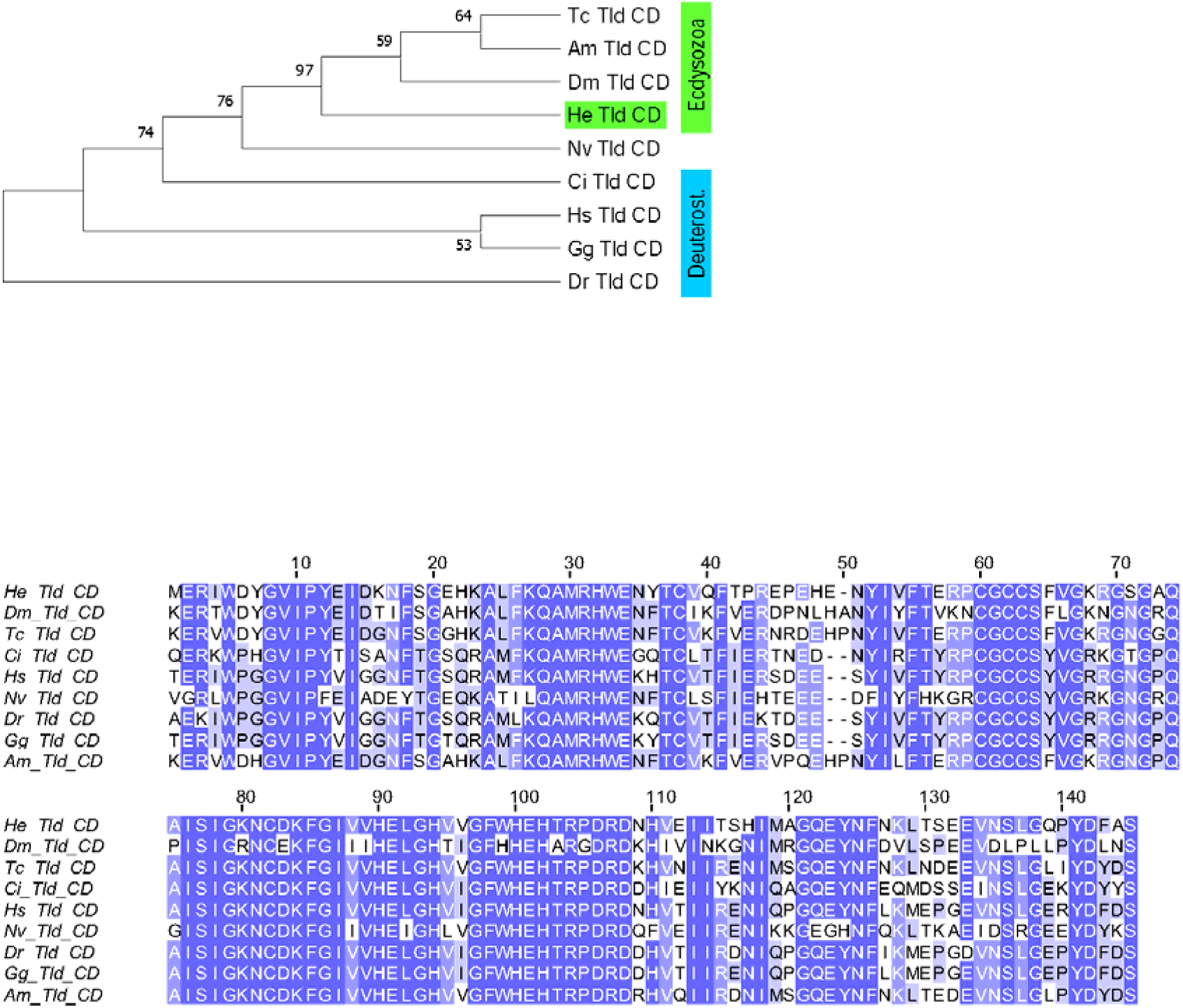
Identification of a Tolloid homolog in *H. exemplaris*. Above: Maximum Likelihood phylogenetic reconstruction of Tolloid and related amino acid sequences. Branch support out of 100 is given at each node. *H. exemplaris* sequences are highlighted. Protein families are indicated by colored bars to the right of the tree. Deuterost = Deuterostomes. Below: Alignment used for phylogenetic reconstruction from conserved domain among Tolloid homologs from several species. Genes include 2-letter prefixes indicating species listed in Table 1.

**Supplementary Figure 8.**
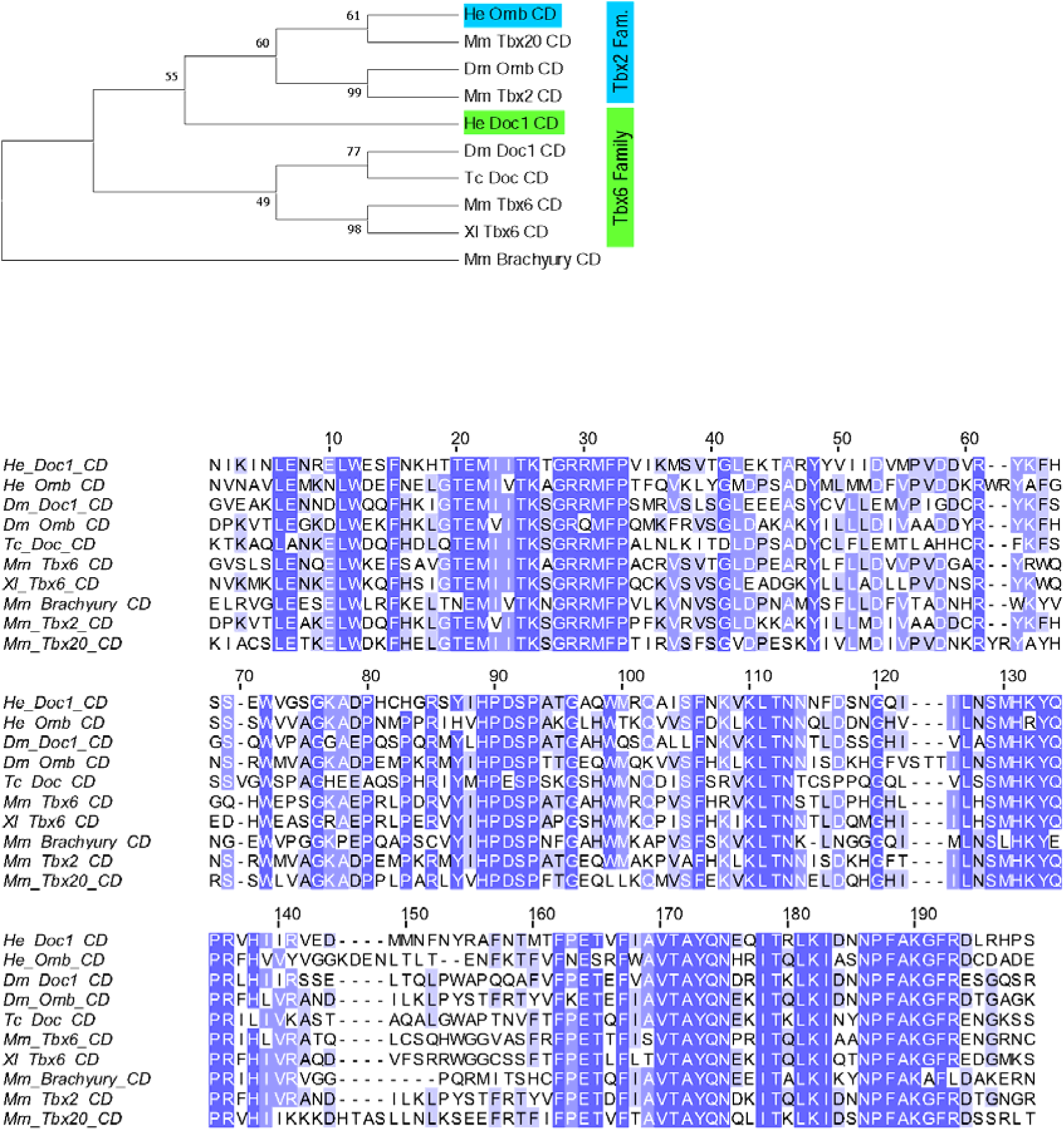
Identification of a Doc1 homolog in *H. exemplaris*. Above: Maximum Likelihood phylogenetic reconstruction of Dorsocross (Doc) and Optomotor Blind (Omb) and related amino acid sequences. Branch support out of 100 is given at each node. *H. exemplaris* sequences are highlighted. Protein families are indicated by colored bars to the right of the tree. Below: Alignment used for phylogenetic reconstruction from conserved domain among Dorsocross homologs from several species. Genes include 2-letter prefixes indicating species listed in Table 1.

**Supplementary Figure 9.**
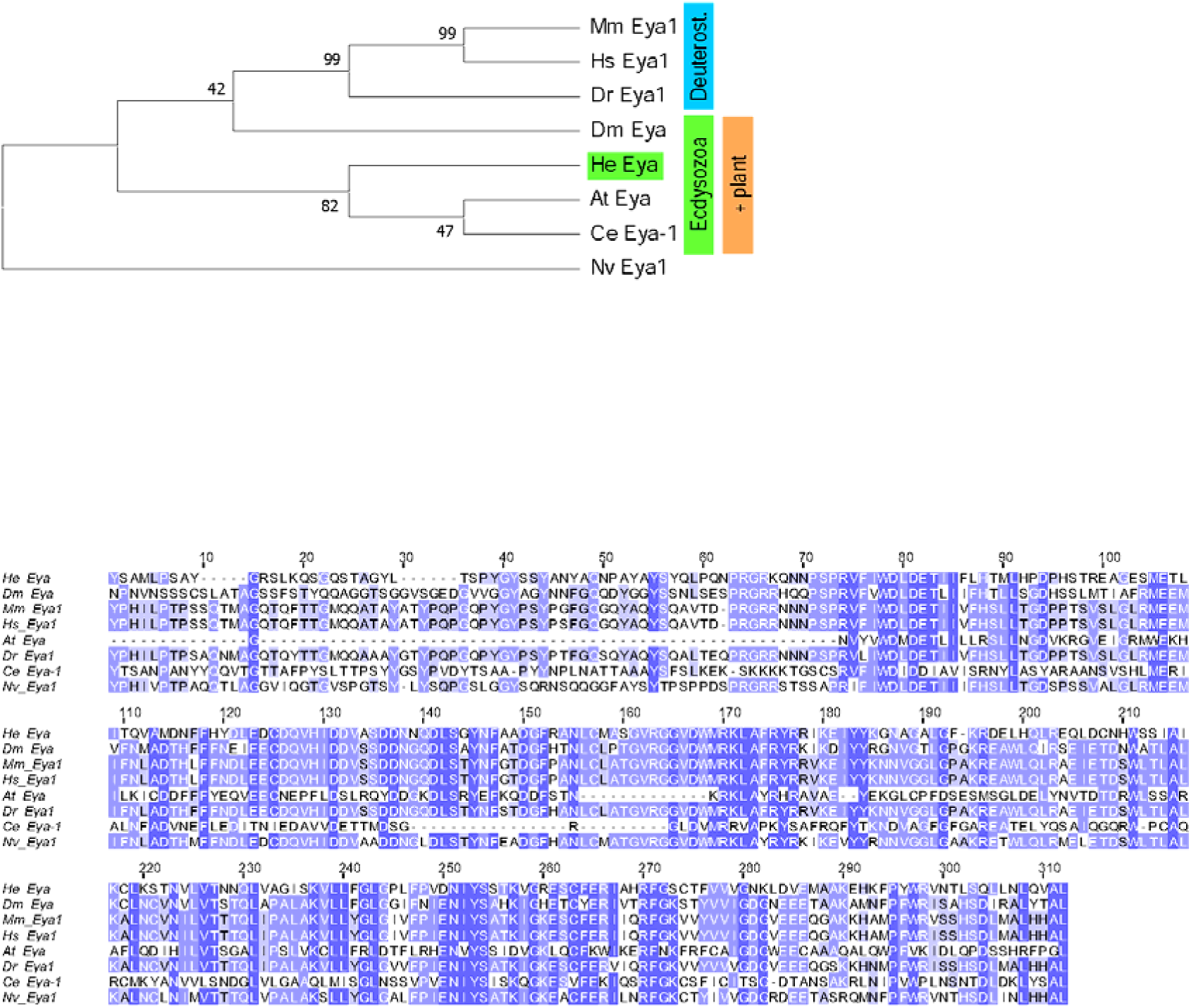
Identification of an Eya homolog in *H. exemplaris*. Above: Maximum Likelihood phylogenetic reconstruction of Eyes absent (Eya) and related amino acid sequences. Branch support out of 100 is given at each node. *H. exemplaris* sequences are highlighted. Protein families are indicated by colored bars to the right of the tree. Deuterost = Deuterostomes. Below: Alignment used for phylogenetic reconstruction from conserved region among Eya homologs from several species. Genes include 2-letter prefixes indicating species listed in Table 1.

**Supplementary Figure 10.**
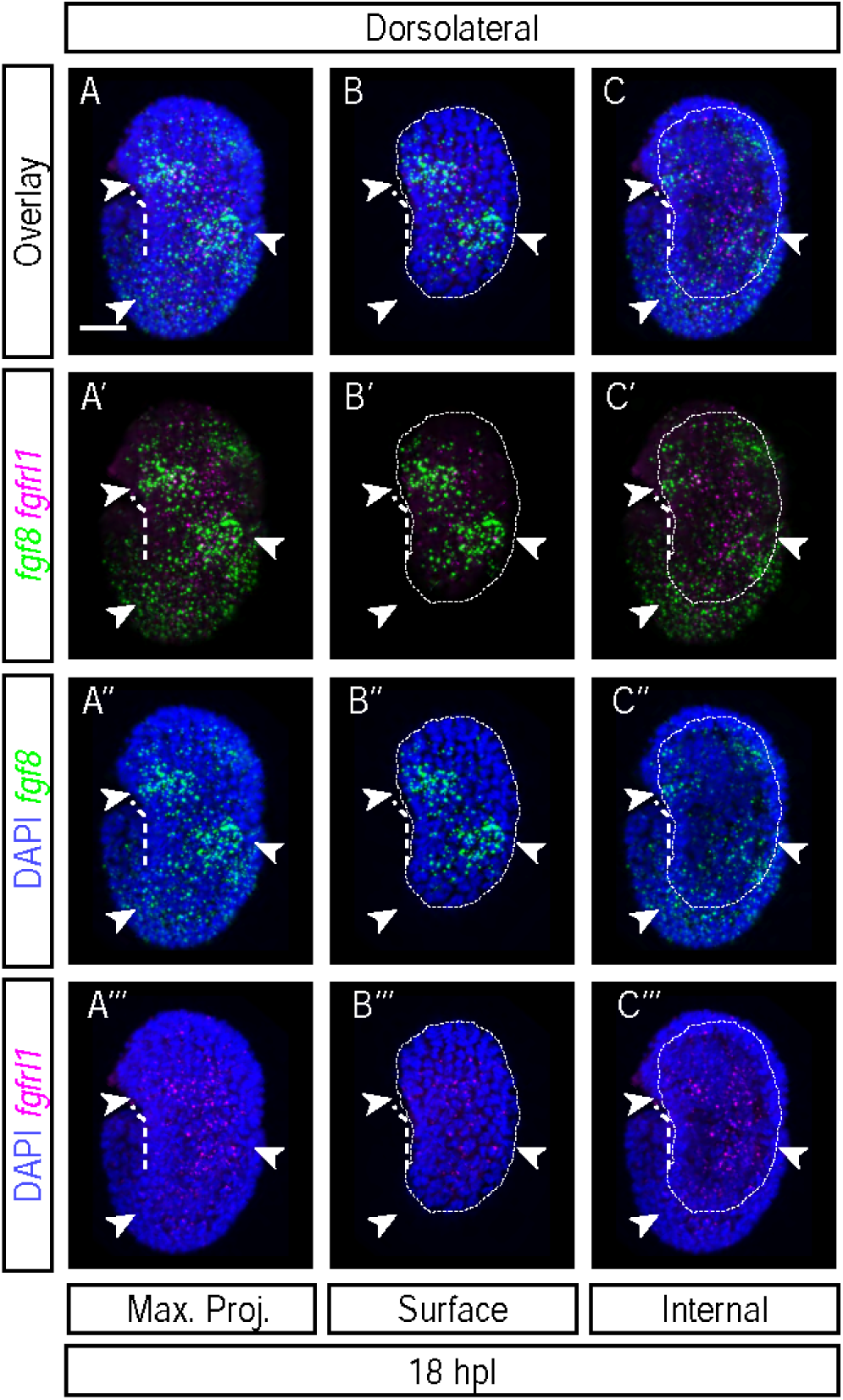
Expression patterns of *fgf8* and *fgfrl1* mRNAs at elongation stage (18 hpl). A-C. Enrichment patterns of *fgf8* and *fgfrl1* mRNAs at 18 hpl (n=2 experiments, 8 and 2 embryos). A. Maximum intensity projection of embryo from a dorsolateral view. B. Projection of uppermost layers of the embryo in A. C. Projection of internal layers of the embryo in A. Region outlined with a dashed line indicates the region with signal in the upper layers projected in B, which are omitted in the internal layers projected in C. Arrowheads indicate ectodermal patches of *fgf8* signal. Left dorsolateral view. Scale bar = 10 μm.

**Supplementary Figure 11.**
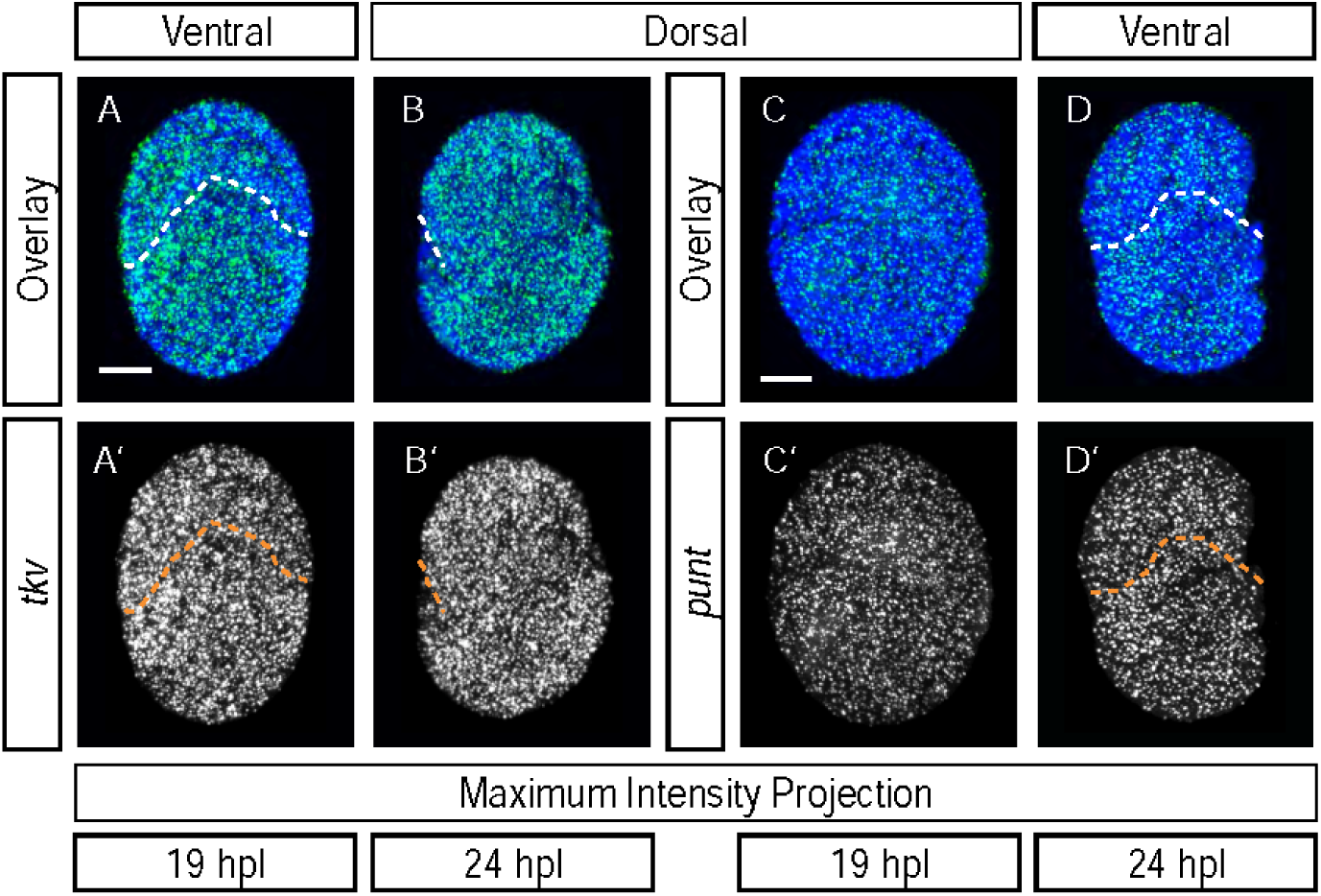
Expression patterns of *tkv* and *punt* at ectodermal segmentation stage (24 hpl). A-B. Enrichment pattern of *tkv* mRNAs at 19 hpl (A, n=1 experiment, 6 embryos) and 24 hpl (B, n=2 experiments, 7 and 4 embryos) by FISH. C-D. Enrichment pattern of *punt* mRNAs at 19 hpl (A, n=1 experiment, 4 embryos) and 24 hpl (B, n=2 experiments, 4 and 4 embryos) by FISH. All images are maximum intensity projections. Embryos are all in ventral or dorsal view, as indicated. Scale bar = 10 μm.

**Supplementary Figure 12.**
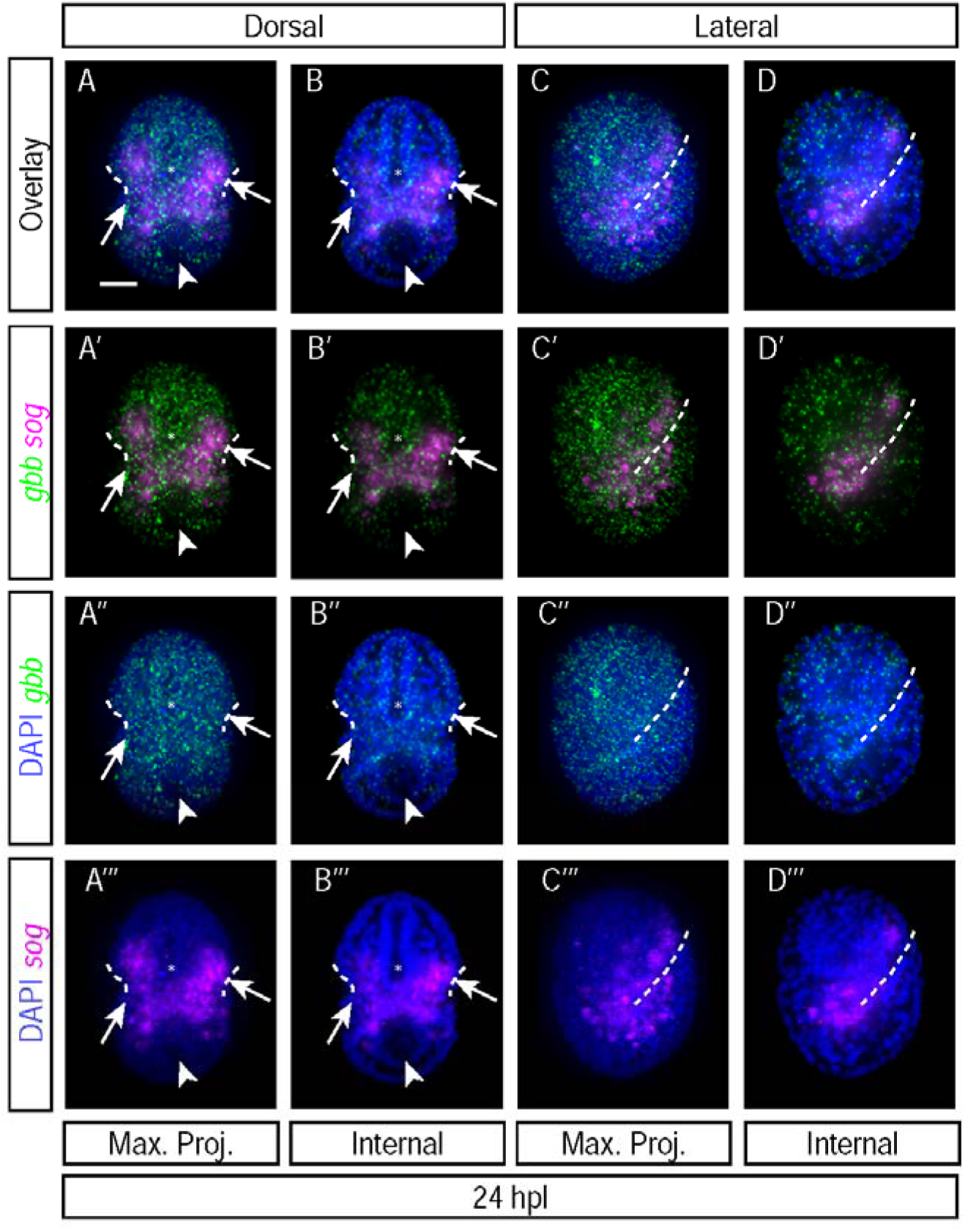
Expression patterns of *gbb* and *sog* at ectodermal segmentation stage (24 hpl). A-D. Enrichment patterns of *gbb* and *sog* mRNAs at 24 hpl (n=1 experiment, 11 embryos). A and C Maximum intensity projections of embryos from a dorsal (A) and lateral (C) view. B. Projection of internal layers of the embryo in A. D. Projection of internal layers of the embryo in C. Arrowheads in A-B indicate region of the PGCs, which is not enriched for *gbb*. Arrows in A-B indicate bands of cells enriched for *sog* on either side of the developing mouth. Asterisks in A-B indicate the hollow space within the developing foregut, which is not enriched for *sog* in its epithelium, but which does contain *gbb* signal. Dorsal view in A-B. Right lateral view in C-D. Scale bar = 10 μm.

## Notes

### Competing Interest Statement

The authors have declared no competing interest.

### Summary of Updates

Author list updated; manuscript structure revised to combine results and discussion in narrative form; discussion augmented to include new interpretation of expression patterns and new discussion points

### Funding

- National Science Foundation: IOS 2028860
- National Institutes of Health: T32GM135128
- UNC Chapel Hill Royster Society of Fellows

